# Metformin Intervention Prevents Cardiac Dysfunction in a murine model of Adult Congenital Heart Disease

**DOI:** 10.1101/396416

**Authors:** Julia C. Wilmanns, Raghav Pandey, Olivia Hon, Anjana Chandran, Jan M. Schilling, Qizhu Wu, Gael Cagnone, Preeti Bais, Vivek Phillip, Heidi Kocalis, Stuart K. Archer, James T. Pearson, Mirana Ramialison, Joerg Heineke, Hemal H. Patel, Nadia A. Rosenthal, Milena B. Furtado, Mauro W. Costa

## Abstract

Congenital heart disease (CHD) is the most frequent birth defect worldwide and the number of adult patients with CHD, now referred to as ACHD, is increasing. However the mechanisms whereby ACHD predisposes patients to heart dysfunction are still unclear. ACHD is strongly associated with metabolic syndrome, but how ACHD interacts with poor modern lifestyle choices and other comorbidities, such as hypertension, obesity and diabetes, is mostly unknown. Using a genetic mouse model of ACHD we showed that ACHD mice placed under metabolic stress (high fat diet) displayed decreased heart function. Comprehensive physiological, biochemical and molecular analysis showed that ACHD hearts exhibited early changes in energy metabolism that preceded cardiac dysfunction. Restoration of metabolic balance by metformin prevented the development of heart dysfunction in ACHD mice. This study reveals that early metabolic impairment reinforces heart dysfunction in ACHD predisposed individuals and diet or pharmacological interventions can be used to modulate heart function and attenuate heart failure and may be an important avenue for intervention in ACHD.

## Introduction

Successful corrective interventions for CHD malformations have led to improved patient survival into adulthood, causing a staggering 60% increase in patients presenting with adult congenital heart disease (ACHD) (Alshawabkeh & Opotowsky, 2016; Marelli et al, 2014). Consequently, the number of ACHD genetically predisposed individuals is on the rise, affecting approximately 2.4 million patients in the US alone (Marelli et al, 2014). Although early corrective interventions for CHD are successful, CHD patients have a higher risk to develop progressive cardiac dysfunction during adulthood. Among the ACHD population the prevalence of comorbidities such as renal, pulmonary, hepatic and vascular dysfunctions, as well as psychiatric diagnosis are elevated. These major health risks drastically increase the morbidity and mortality of ACHD patients (Lui et al, 2017). The high prevalence of associated metabolic disorders in ACHC is also merits further investigation, given the multiple prophylactic options currently available in the clinic.

The continuous decline in physical activity and increased caloric intake has resulted in an obesity epidemic in developed countries as the major health risk of the 21^st^ century. It is estimated that approximately 63.3% of the American population is currently overweight or obese (Writing Group et al, 2016). ACHD intrinsic congenital heart dysfunction is exacerbated by external environmental factors, including obesity, diabetes and/or high blood pressure, all of which can trigger heart failure (Islam et al, 2016; Roche & Silversides, 2013). It is therefore imperative to understand why today’s lifestyle set ACHD patients at an even higher, life-threating risk than the general population. Despite the existence of large epidemiological datasets, not much is currently known about how ACHD predisposes patients to heart failure upon metabolic stress. The present study focuses on how obesity affects cardiac function in our recently characterized ACHD model (Dietz, 2015; Ogden et al, 2014). Using genetically predisposed mice (Furtado et al, 2017) and diet as a cardiac stressor, we describe a preexistent imbalance in the metabolic state of ACHD hearts. Development of obesity increases the severity of heart dysfunction. This interaction between genetic and metabolic factors ultimately leads to the clinical presentation of heart failure in ACHD. Modulation of energy utilization by Metformin, a drug widely used to treat type 2 diabetes, prevents cardiac dysfunction in ACHD/obesity model and could therefore be considered a preventive intervention for heart failure in ACHD. Our observations also suggest that ACHD is a complex, multifactorial disease that can be modulated by changes in global metabolism. Patients at risk for ACHD show intrinsic metabolic dysfunction that is aggravated by exposure to a modern lifestyle environment leading to cardiac dysfunction, and therefore should be differentially monitored.

## Methods

### Mouse lines

*Nkx2-5*^*C/+*^ *and Nkx2-5*^*183P/+*^ (ACHD) mice have been previously characterized (Furtado et al, 2017). All mice were maintained on a C57BL/6J background, housed at Monash Animal Services, Australia or at The Jackson Laboratory, USA. This investigation conforms with the Guide for the Care and Use of Laboratory Animals published by the US National Institutes of Health (NIH Publication No. 85–23, revised 1996) and requirements under the ethics application MARP-2011-175 (Monash University) and ACUC 16011 (The Jackson Laboratory). Control mice used were either *Nkx2-5*^*C/+*^ (homologous recombination replacing the I residue with itself; i.e. control = C) or wild-type (WT). No significant molecular or physiological differences were observed between these two control groups (Furtado et al, 2017). Whenever possible, control littermates were used for the experiments. All experiments were performed on adult males.

### Feeding regimen

Six-week-old heterozygous male mice were subdivided into four experimental groups according to feeding regimen (control Low Fat (LF), control High Fat (HF); ACHD LF and ACHD HF). Throughout the study, two animal cohorts were studied: short (weeks 6 to 15) and long (weeks 6 to 30) feeding regimens. All mice were single housed under normal environmental conditions (21.5 ± 1°C with a light-dark cycle of 12hr:12hr). For metformin studies 4 siblings male mice were housed together. Food and water were accessible ad libitum. Low Fat diet (SF13-081/ D12451; total calculated digestible energy from lipids: 12.0%, total calculated digestible energy from protein: 25.8%) and High Fat diet (SF04-001; total calculated digestible energy from lipids: 43.0%, total calculated digestible energy from protein: 21.0%) were purchased from Specialty Feeds Pty Ltd. (Australia) and Research Diets Inc. (USA). Metformin at the concentration of 5 mg/mL was added to the water from 8 weeks of age until the end of the protocol *at libidum*. Weight and food consumption were recorded weekly at 10a.m ± 2 hr from the age of 6 weeks until week 15 for short feeding regimen groups and further calculated at week 30 for long feeding regimen groups.

### Histology

Mice were euthanized via CO_2_ asphyxiation, perfused with HBSS (Thermo), followed by heart dissection. Hearts were further incubated for 5 minutes in 60mM KCl for synchronization of heart cycle before fixation. Tissues were fixed in 4% PFA at 4°C overnight and further processed for paraffin sectioning following standard dehydration and embedding protocols. Adult thyroid samples were cut with a 14µm thickness, adult hearts with a 10µm thickness. The Masson’s Trichrome staining was performed by the Monash Histology Facility. For myocyte area measurements, transverse sections were stained with Wheat Germ Agglutinin Alexafluor 488 conjugate (WGA, Thermo). All images were obtained using Olympus DotSlide (Japan). Mitochondrial density was analyzed by immunofluorescence on heart cryosections using TOMM20 rabbit polyclonal antibody (Abcam) raised against the *Translocase of Outer Protein Membrane 20* gene (*Tomm20*). Confocal Z-stack images were performed using Leica SP8. Myocyte area and fluorescent intensity was measured using FIJI software with three replicates for each genotype. For TEM, 50 mg of cardiac tissue as processed by the JAX histology core and sections were evaluated using JM-1230 microscope (JEOL) coupled to AMT 2K camera (Advanced Microscopy Techniques). Statistical significance of data was determined by ANOVA or Student’s t-test.

### Magnetic Resonance Imaging (MRI), Echocardiography and ECG analysis of cardiac function

15- and 30-week old adult mice were anaesthetized with 2% ± 0.2% isoflurane for non-invasive imaging in a prone position. For MRI, imaging was performed in spontaneously breathing mice in a 9.4 Tesla MRI scanner (Agilent, USA) to assess left ventricular and right ventricular chamber dimensions in systole and diastole using T1 and T2-weighted anatomical sequences. Breathing was also monitored to avoid measurement distortions during the breathing cycle. Measurements of cardiac function were performed using Segment software (Heiberg et al, 2010) analysed blinded to genotype by two independent researchers. Echocardiogram analysis was performed using Vevo 2100 (FUJIFILMS Visualsonics). ECG was recorded in anaesthetised mice using Powerlab data acquisitions system (ADInstruments, USA) and analysed using LabChart8 software (ADInstruments, USA).

### Metabolic Measurements

For accurate measurement of weight and body fat composition we performed dual-energy X-ray absorptiometry (DeXa) using Lunar PIXImus Densitometer (GE Medical Systems, USA) or Echo MRI instrument. For DeXa analysis, mice were anaesthetized with 1.7% ± 0.2% isoflurane. Whole-body metabolic changes activity and energy expenditure was assessed using metabolic cages (Comprehensive Cage Monitoring System – Promethion). Mice were maintained on a 12hr light:dark cycle with access to food and water *ad libitum* in cages undisturbed for 120 hours to collect CO_2_ production, O_2_ consumption, locomotor activity, body mass, food and water intake. Voluntary exercise capacity in running wheels (Med Associates, USA) was measured by recording running distance and circadian activity for 5 continuous days. Metabolite levels of cholesterol, triglycerides, glucose and NEFA were quantified from mouse sera using the Clinical Assessment Services, The Jackson Laboratory.

### Adult cellular respiration analysis

Mitochondrial function in mechanically permeabilized fibers from the left ventricular free wall was evaluated using the OROBOROS O2k Respirometer added to the oxymetry chamber containg MiR06 media with 25µM Blebbistatin added (EGTA 0.5 mM, MgCl_2_ 6H_2_O 3mM, Lactobionic acid 60mM, Taurine 20mM, KH_2_PO_4_ 10mM, HEPES 20mM, D-Sucrose 110mM, BSA 1g/l, Catalase 280 U/ml, pH 7.1, at 30 °C) (Gnaiger et al, 2000). Fatty acid oxidation and Complex I protocol were adapted from a previously published protocol(Schopf et al, 2016) depicted in **Figure 4E**.

### Metabolomics and western blot analysis

Unbiased discovery metabolite platform was performed at West Coast Metabolomics Center at UC Davies, USA. Biventricular apex fragments were snap frozen in N_2_ and shipped to the center in dry ice. The metabolomics platform used consisted of four different combinations of liquid chromatography (LC) and mass spectrometry ionization using Agilent 6890 GC followed by Pegasus III TOF MS. Relative abundance of 273 unique metabolic features (109 metabolites of known structures and 164 unique features of unknown structure), normalized to internal standards were measured at each time point. Metaboanalyst 3.0 software (Xia & Wishart, 2016) was used for bioinformatics analysis. Interquartile range based filtering was used to remove features unlikely to be used for modelling purposes(Hackstadt & Hess, 2009). The data was log transformed and autoscaled (Dieterle et al, 2006). Fold Change (FC) analysis (before column normalization), t-tests, and volcano plot were generated comparing the mutant samples with the control samples at each time point. Protein changes were analysed by western blot analysis using proteins extracted from biventricular fragments as previously described (Furtado et al, 2017). Antibodies used are described in **Supplementary Table 4**.

### Functional annotation enrichment analysis of Nkx2-5 ChIP peak regions

Publicly available chromatin immunoprecipitation followed by deep-sequencing (ChIP-seq) data (GSE35151) (van den Boogaard et al, 2012) was re-analyzed by our group. Unsupervised enrichment analysis of genome-wide occupancy profile of Nkx2-5 in adult mouse whole heart was performed using Genomic Regions Enrichment of Annotations Tool (McLean et al, 2010). We performed gene ontology (GO) analysis using GREAT default setting (5 kb proximal + 1 kb basal, up to 1 Mb distal, mm9 reference genome) calculating statistical enrichments by associating genomic regions with nearby genes and applying the gene annotations to the regions.

### RNAseq, Bioinformatics and validation

Ventricular heart samples from 15-week old adult males of each genotype (*Nkx2-5*^*183C/+*^, WT and *Nkx2-5*^*183P/+*^ mice) on a high fat diet were used as biological replicates for RNA sequencing. Total RNA extraction was performed using the mirVana kit (Thermo, USA) and cleared of genomic DNA contamination using in column DNAse treatment (Qiagen, Germany). Samples were further processed by the Medical Genomics Facility at Monash University. RNA-seq data (paired-end reads of 100 nt) was trimmed for quality and adapter contamination using Cutadapt (Martin, 2011) as implemented in Trim Galore (Phred quality threshold of 28, minimum adapter overlap 3 nt, adaptor sequence AGATCGGAAGAGC). It was then mapped to the mouse genome (mm10) using STARv2.4.0(Dobin et al, 2013), and reads were tallied within gene features using HTSeq (Anders et al, 2015). From the resulting read-count matrix, genes with more than 10 reads in any sample were kept and differentially expressed genes were called using limma.voom (Law et al, 2014). Targets with a FDR cut-off of 0.1 were further selected for validation using qPCR. Ingenuity Pathway Analysis (IPA) software (Qiagen, Germany) was used to analyze differentially regulated pathways and functions in datasets. cDNA synthesis was performed in the same samples using Superscript VILO (Thermo) followed by SYBR green reactions (Roche, USA) and analysed using the LightCycler480 (Roche, USA). Relative quantification was obtained using *Hprt* as a normalizer. Primers used are described in **Supplementary Table 4**.

### Statistics

Data as presented as a mean ± SEM. Statistical significance of data was determined by ANOVA and Student’s t-test. P values < 0.05 were considered as significant.

## Results

### ACHD mice show increased susceptibility to weight gain and development of obesity

In a previous study, we described a point mutation in the *NKX2-5* gene found in a familial ACHD cohort. We engineered the same human mutation in mice and showed that our model reproduced all clinical features of the disease (Furtado et al, 2017). To explore the interplay between ACHD, metabolism and heart function, we subjected ACHD mice to control conditions under low fat (LF) and high fat (HF) diet as a metabolic stressor (**Figure 1A**). At the onset of the feeding regimen (6 weeks of age), ACHD mice had significantly lower weight when compared to control mice (**Supplementary Figure 1**). Surprisingly, after 9 weeks of diet, ACHD mice showed a significantly higher weight gain when compared to control mice on HF diet. The increased weight gain was attributed to higher body fat, as lean body weight did not differ between groups (**Figure 1B**). No difference in weight was observed between experimental groups subjected to LF diet. In summary, ACHD mice showed comparatively higher fat accumulation and weight gain when subjected to higher caloric intake, indicating that ACHD and diet have combined additive effects on weight gain and therefore ACHD mice are more predisposed to develop obesity.

**Figure 1.**
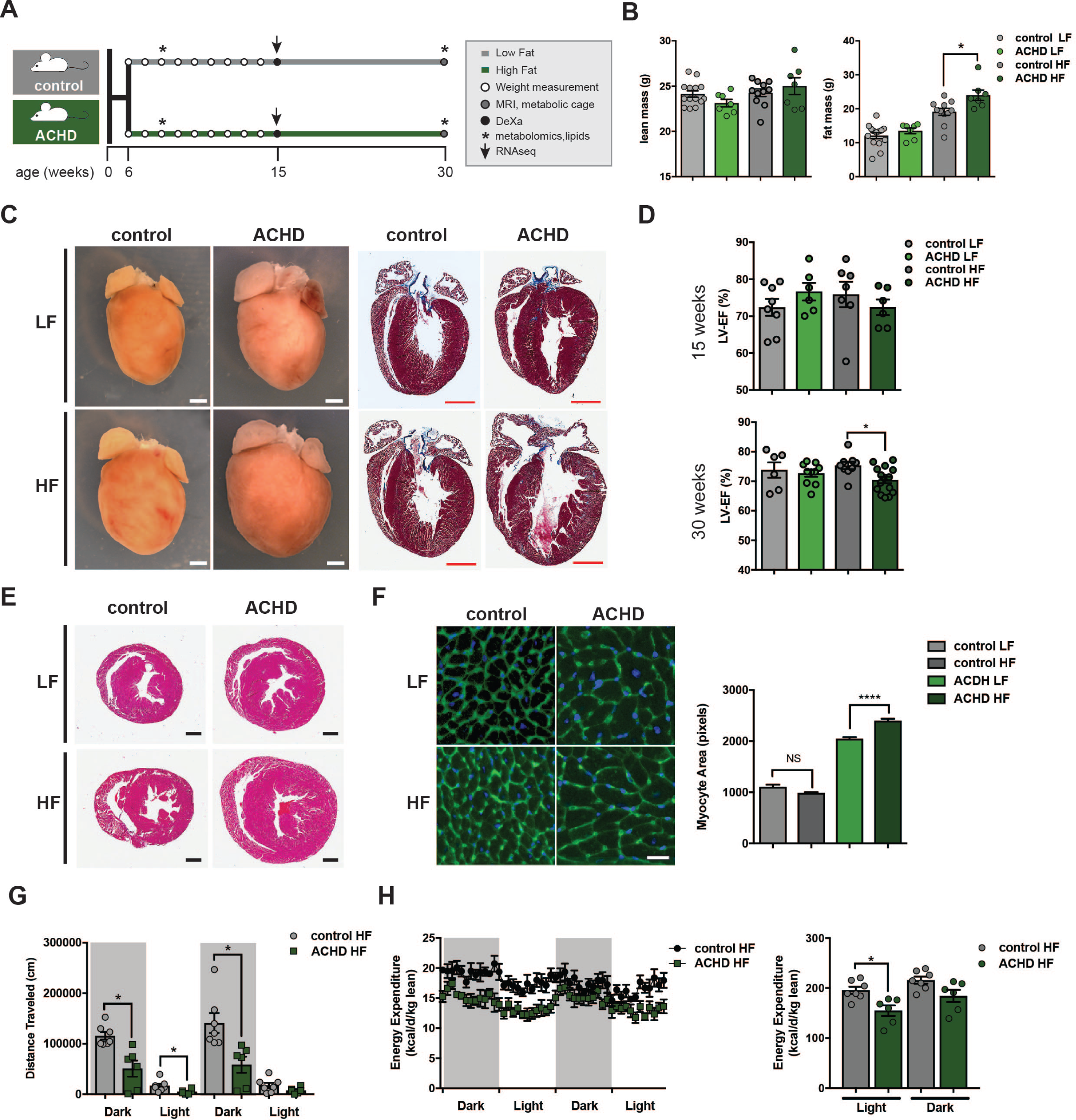
Metabolic stress (HF) diet associated with ACHD leads to cardiac morphological changes and decreased function. (A) Experimental groups and design. (B) Body weight measurements at 30 weeks show that ACHD mice have increased fat mass when compared to controls. (C) Whole-mount and trichrome stained sections at 30 weeks showing increased dilation in ACHD hearts on HF diet when compared to control mice. (D) Functional impairment detected as decreased left ventricle ejection fraction (LV-EF) in 30-week-old ACHD mice when compared to controls. No significant changes were seen at 15 weeks of age. (E-F) Transverse section and quantification of myocyte cross-sectional area of WGA stained samples showed hypertrophy in ACHD hearts that is exacerbated by HF diet. (G-H) ACHD mice display decreased running capacity and energy expenditure, consistent with onset of heart failure (N=6; Mean ± SEM, **p<0.01, Student’s t-test). LF: low-fat.

### Obesity triggers cardiac dysfunction in ACHD

We have previously shown that most disease features developed in the ACHD model are functionally compensated in homeostasis in early adulthood (Furtado et al, 2017), except for RV dysfunction, defined by a significant decrease of right ventricular ejection fraction (EF) (**Figure 1C-D**). Short-term 9-week HF feeding regimen (15 weeks of age) did not significantly affect cardiac function (**Figure 1D**, **Supplementary Figure 2A**). However, the long-term 24-week regimen (30 weeks of age) caused a significant decrease in EF for both right (RV) and left (LV) ventricles, as well as enhanced RV and right atrial (RA) dilation in ACHD mice (**Figure 1C, D; Supplementary Figure 2B).** Once subjected to HF diet, murine ACHD hearts showed increased wall thickness, eccentric ventricular hypertrophy with dilation, and an increase in transverse myocyte area (**Figure 1E-F**). Hypertrophy was also evident in ACHD mice subjected to LF diet but was enhanced on HF diet. No hypertrophy was seen in control mice independent of the diet of choice, demonstrating the importance of genetic predisposition for the hypertrophic response. In summary, ACHD mice showed several morphological abnormalities and cardiac dysfunction prior to the development of Heart Failure with Preserved Ejection Fraction (HFpEF) (Ferrari et al, 2015) when subjected to metabolic stress (HF diet), suggesting that challenging a weakened ACHD heart with metabolic stress accelerates progression to heart failure.

### Obesity triggers global metabolic energy dysfunction in ACHD mice

When provided with voluntary running exercise, ACHD mice on HF diet at 30 weeks displayed significantly reduced physical performance, as measured by decreased running distance (dark cycle; **Figure 1G**). No significant changes were detected in time spent in the wheel, food intake and activity, indicating that ACHD mice were active but not running at the same speed as control mice (**Supplementary Figure 3A, B**). Overall energy expenditure of mice on HF diet was also significantly reduced in ACHD mice (**Figure 1H**), with decreased levels of O_2_ consumption (VO_2_) and CO_2_ production (VCO_2_) rates, while still maintaining normal respiratory quotient (RER). These changes were not observed when ACHD mice were subjected to a Western diet formulation, which contains high sugar (Graham et al, 2016), indicating that cardiac dysfunction was likely mediated by increased availability of fatty acids in the HF diet. Together, these data confirm that nutritional imbalance exacerbates heart dysfunction in ACHD mice.

To determine how global metabolism was affected, we monitored serum levels of glucose and lipids of ACHD mice on HF diet. While no significant changes were seen at 6 weeks, higher plasma levels of triglycerides and a tendency towards increased cholesterol and free fatty acids (NEFA) at 30 weeks was observed (**Figure 2A**). Glucose handling was also affected in ACHD mice on HF diet, with an improved response when compared to control mice (**Figure 2B**).

**Figure 2.**
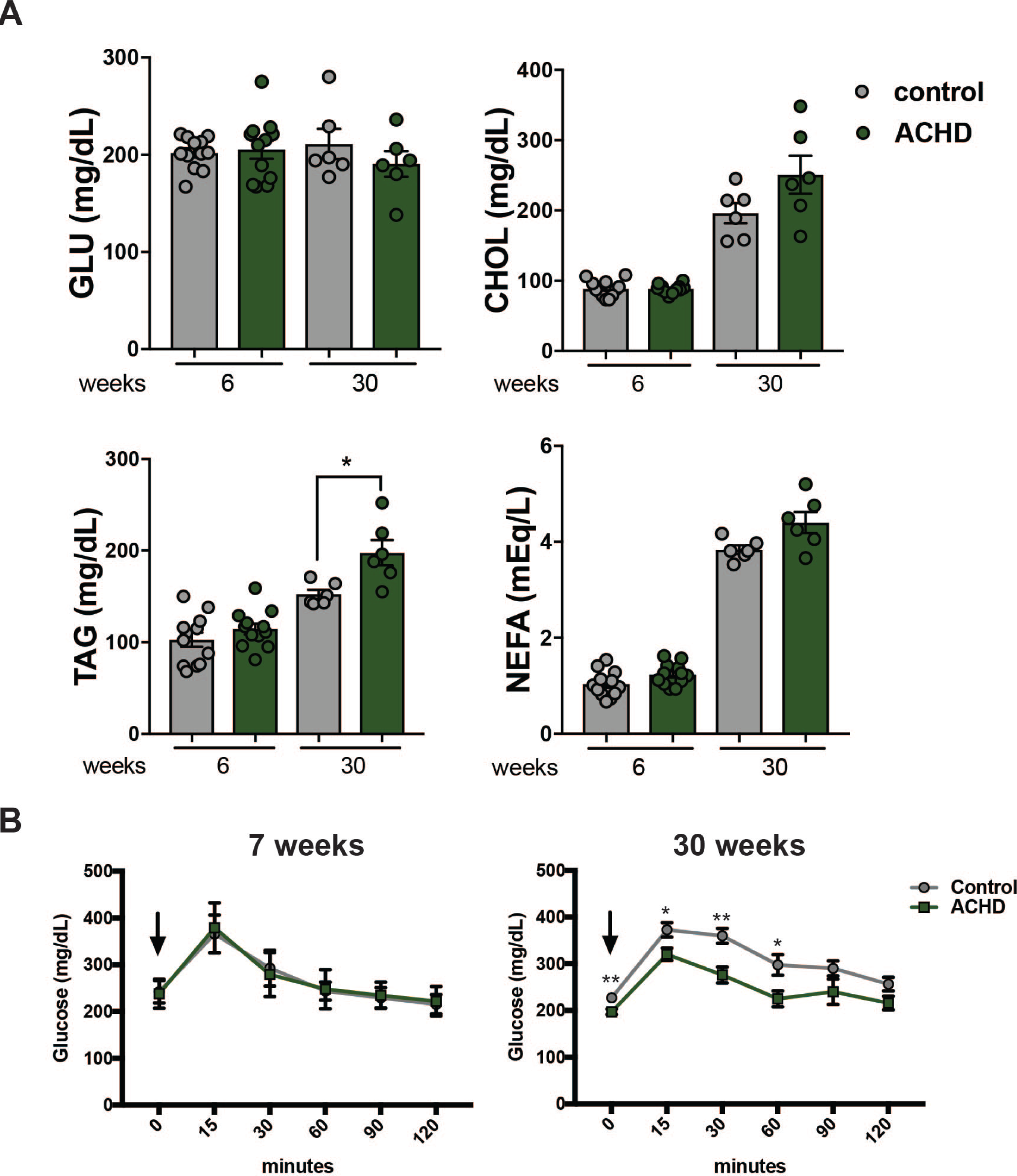
Metabolic disturbances in ACHD hearts. (A) Serum glucose and lipid analyses demonstrate small increases in cholesterol and free fatty acids and a significant increase in triglycerides in ACHD mice at 30 weeks. No significant changes were seen at 6 weeks of age. Blood glucose levels show a small non-significant decrease in ACHD mice. (B) ACHD mice display improved glucose handling when compared to controls at 30 weeks of age during glucose tolerance test, while no changes were observed at 7 weeks. (Mean ± SEM, *p<0.05, **p<0.01; Student’s t-test). GLU-glucose; CHOL-cholesterol; TAG-triglycerides; NEFA-nonsterified free fatty acid.

### Primary heart metabolic dysfunction in ACHD mice is driven by changes in energetic substrates

*Nkx2-*5 is a transcription factor essential for heart formation during embryonic development, although the molecular consequences of its dysfunction in adult hearts are still largely unknown. Analysis of chromatin immunoprecipitation followed by deep-sequencing (ChIP-seq) of adult hearts using gene ontology (GO) and molecular signature database (MSigDB) revealed that the human NKX2-5 protein binds the promoter regions of essential genes associated with cellular metabolism and energy production (**Figure 3A, Supplementary Table 1**), a previously unrecognized phenomenon. To further explore the molecular mechanisms underlying the primary metabolic alterations in ACHD hearts, we performed transcriptome analysis (RNAseq) on short-term HF-fed mice, before cardiac dysfunction was established. This analysis focused on investigation of genes that were differentially expressed in obese control and ACHD mice, to expose molecular differences of the metabolic stress model (ACHD + obesity). Relative quantification showed 1264 genes differentially expressed between control and ACHD obese hearts. Ingenuity Pathway Analysis (Qiagen) revealed 41 predicted upstream regulators of affected genes **(Figure 3B)**, including cardiac transcription factors known to partner with NKX2-5, four of which are involved in cardiogenesis and cardiac function. The canonical pathway of cardiac hypertrophy was significantly changed in ACHD (z-score=-1.347; p-value=0.00011), confirming our finding of eccentric hypertrophy (**Figure 1E, F**).

**Figure 3.**
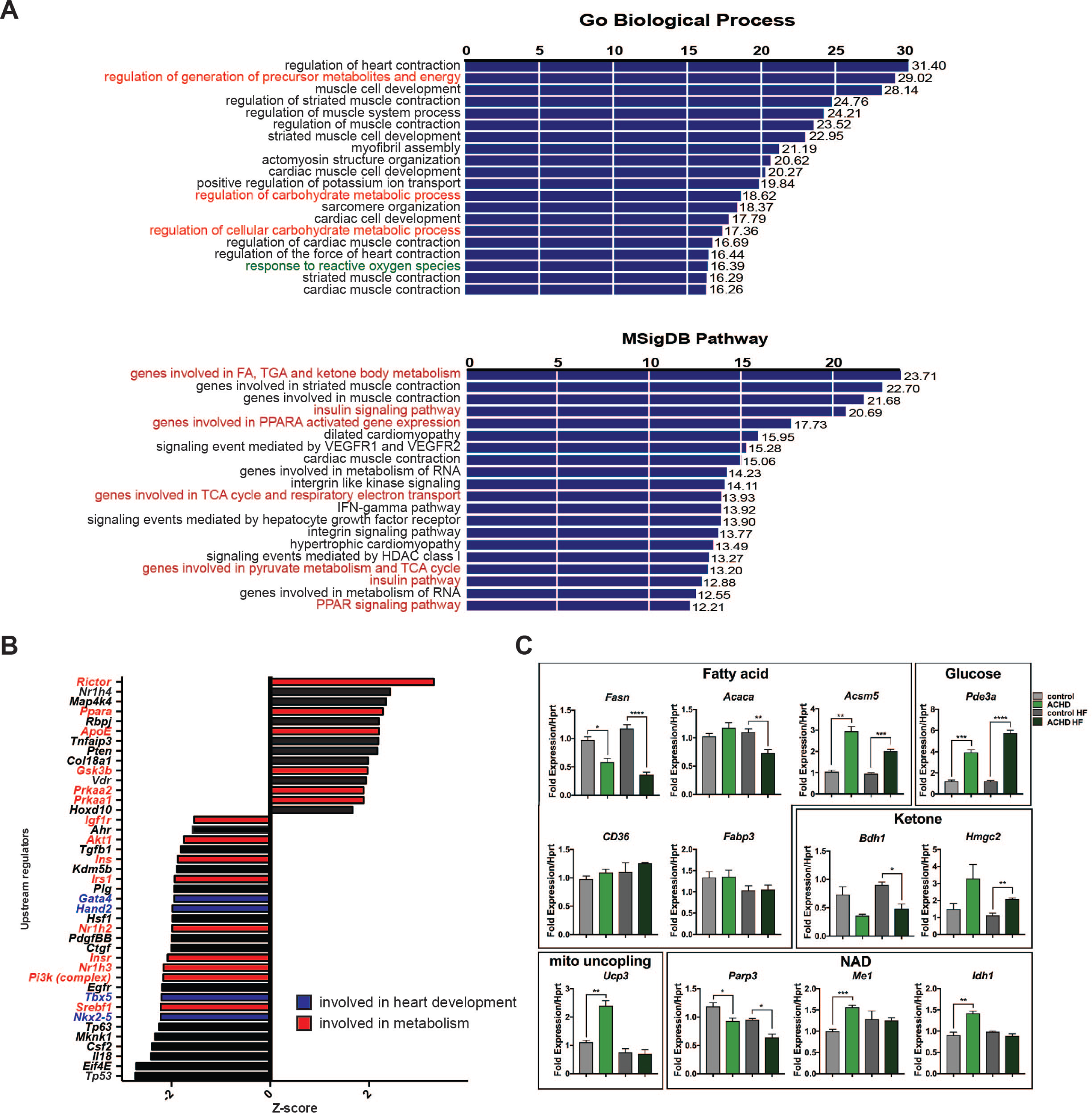
NKX2-5 regulates energy handling in adult hearts. (A) Data mining of ChIP-seq against NKX2-5 in adult hearts (GEO GSE5151) shows high prevalence of genes involved in metabolism (red) in both Go Biological Process and MSigDB pathways. (B). Ingenuity Pathway Analysis (IPA) shows increased prevalence of predicted upstream regulators for metabolism (red) and heart development (blue) processes using RNAseq data of ACHD obese hearts. (C) qPCR analysis confirms metabolic changes associated with energy flexibility in ACHD/obesity model at 15 weeks under normal chow or HF diet. Data presented as fold expression compared to control samples normalized by *Hprt* levels. (N=3; Mean ± SEM, *p<0.05, **p<0.01; ***p<0.001, ****p<0.0001, Student’s t-test).

Among genes involved in the hypertrophic response, *Rock* and *Mlc* complexes and a large subset of other effector genes such as *Gata4, Hand2, Mef2* and *p300* were up regulated in ACHD mutants (**Supplementary Figure 4**). These genes trigger an increase in contractility, leading to long-term cardiac hypertrophy (Dai et al, 2002; Dirkx et al, 2013; Hartmann et al, 2015; Heineke & Molkentin, 2006; Yanazume et al, 2003). Furthermore, *eIF4E* (mRNA cap-binding protein) and *Mlnk1* (eIF4E kinase), known essential regulators of cell size control and global protein synthesis, were also affected in ACHD hearts. Interestingly, 37% of upstream regulators were associated with metabolic pathways and lipid processing, including *Srebf1, Nr1h2-3, Insr*, and *ApoE*, among others. This analysis corroborates the global whole-body metabolic data, indicating that disturbances in heart metabolism precede progression to heart dysfunction in ACHD.

Another affected pathway was PPAR signalling, where *Ppara, Prkaa1* and *Prkaa2* were highlighted as potential regulators in ACHD mice. These proteins are key components in fatty acid metabolism and are normally activated to compensate for energy deprivation, which is a common feature of heart failure (Djouadi et al, 1998; Luptak et al, 2005) (**Figure 3B**). Transcriptional changes directly linked to metabolism were further confirmed by qPCR (**Figure 3C**). In particular, decreased levels of *Fasn* and *Acaca* and increased levels of *Acsm5* indicated marked reduction of fatty acid synthesis and increased fatty acid oxidation, presumably via loss of inhibition of CPT1 (Lopaschuk et al, 2010), whereas *Fabp3* and *CD36* fatty acid receptor transcripts were not changed, suggesting no alteration in fatty acid uptake. In agreement with these observations, increased levels of *Pde3b* suggested enhanced glucose metabolism, although no significant changes were seen in *Insr* expression. Ketone body metabolism gene *Bdh1* showed decreased levels, while *Hmgcs2* was increased (Aubert et al, 2016).

Genes associated with mitochondrial uncoupling/protection (*Ucp3*) and NAD+ metabolism (*Me1, Idh1* and *Parp3*) were also affected. Most of these genes are normally activated in heart failure (Luptak et al, 2005). These changes in gene expression are likely associated with the shift in energy signature (Doenst et al, 2013; Ventura-Clapier et al, 2004), indicating that ACHD mice show imbalanced energy utilization, and confirm that metabolic changes in ACHD mice are enhanced by HF diet and precede heart failure.

### Imbalanced cardiomyocyte energy handling capacity precedes heart dysfunction in ACHD

To further characterize molecular mechanisms associated with ACHD onset of progression to heart failure, we performed a comprehensive metabolic and physiological analysis in early adulthood, before ACHD mice develop cardiac dysfunction. This allowed us to filter out changes that are not directly caused by cardiac dysfunction. At 8 weeks of age, ACHD mice showed a small but significant decrease in body weight compared to control wild-type mice, associated with decreased lean mass (**Supplementary Figure 1**), although no changes were observed in energy expenditure, VCO_2_/VO_2_ consumption or voluntary exercise capacity (**Supplementary Figure 5A-C**). No significant changes were observed in cardiac electrical activity (**Supplementary Figure 6**), other than increased spread of the QRS interval, consistent with the well-established role for NKX2-5 in the maintenance of the conduction system in mouse models and patients (Chowdhury et al, 2015; Gutierrez-Roelens et al, 2002; Jay et al, 2004; Pashmforoush et al, 2004). Echocardiographic (Echo) analysis of heart function showed small changes in left ventricular function but normal ejection fraction, indicating that imbalance of NKX2-5 protein activity can be compensated for to ensure adequate heart function in early adulthood (**Supplementary Table 2**).

We have previously observed decreased mitochondrial respiratory capacity in neonatal ACHD cardiomyocytes (Furtado et al, 2017). Therefore, we investigated cellular respiration and energy handling in young ACHD hearts at 8-10 weeks of age. Mitochondrial morphology was not changed in ACHD cardiomyocytes compared with controls (**Figure 4A**), but a significant decrease in mitochondrial density was observed (**Figure 4B-D**). In agreement with the data obtained in neonatal cardiomyocytes (Furtado et al, 2017), mitochondrial function in left ventricular free wall fibers of the young adult heart revealed tendency to reduced oxygen flux and fatty acid oxidative capacity in ACHD hearts (**Figure 4E**). Unbiased global metabolomics (untargeted GCMS analysis) showed a significant increase in glycolysis and a small decrease in ketone and fatty acid oxidation (FAO) metabolites in ACHD hearts (**Figure 4F, G** and **Supplementary Table 3**), in particular glucose-1P, glucose 6-P, fructose 6-P for glycolysis and 3-hydroxybutyric acid for ketone metabolism. Furthermore, AMP levels were increased in ACHD hearts but this alteration did not trigger differential AMPK activation (**Figure 4G, H**), suggestive of failure to respond to energetic imbalance in mutant hearts. Metabolomic changes were further confirmed by western blot of heart samples, which revealed a significant increase in PDH (Glucose metabolism), small decrease in CPT1 (Fatty Acid metabolism) and BDH (Ketone metabolism) (**Figure 4H**). These data suggest a shift from FAO towards increased glycolysis in ACHD myocytes, reminiscent of the fetal energetic program. Increased glucose dependency is an adaptive response to maintain heart function under stress conditions (Aubert et al, 2016; Ritterhoff & Tian, 2017; Taegtmeyer et al, 2016). The observed changes in cardiomyocyte energetic pathways indicate that disturbances in metabolic state precede heart dysfunction in ACHD and therefore represent valuable parameters to measure predisposition to heart failure.

**Figure 4.**
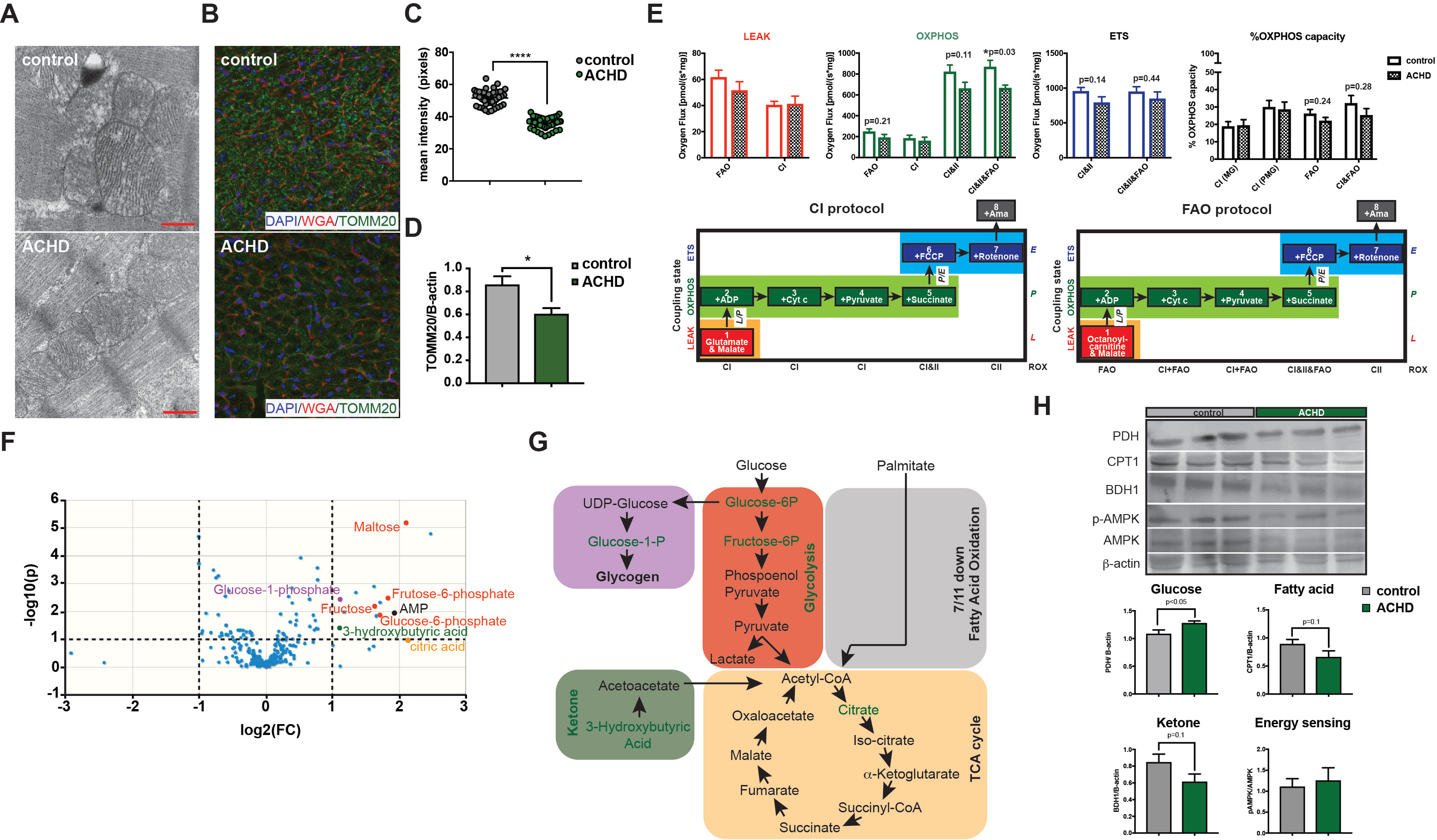
ACHD hearts display early mitochondrial and energy handling defects. (A) Mitochondria of 10-week ACHD hearts show normal morphology but (B-C) decreased density (N=3). (D) TOMM20 protein levels are also decreased in western blot (N=3). (E) No changes in LEAK respiration, decreased oxygen flux under OXPHOS conditions in the presence of ADP, FAO, CI&II, and CI&II&FAO substrates and tendency of decreased maximal respiration of the electron transport system (ETS) are seen in ACHD hearts using the OROBOROS O2k Respirometer oxygen flux. OXPHOS capacity shows a small tendency to decrease in FAO contribution in ACHD hearts, suggesting a FAO driven dysfunction. CI = complex I; CII = complex II; FAO = fatty acid oxidation; M = malate, G = glutamate; P = pyruvate; ROX = residual oxygen flux; (N=6/group). (F) Metabolomics analysis shows over-representation of glycolysis (red dots), ketone (green dot), glycogen (maroon dot), TCA cycle (orang dot) and AMP (black dot) metabolites in ACHD hearts at 8 weeks (N=6). (G) Up-regulated metabolites and processes highlighted in green. (H) Western blot analysis and quantification of enzymes associated with glucose (PDH), FOA (CPT-1), ketone (BDH1) and energy sensing (AMPK)_(N=3). (Mean ± SEM, ****p<0.0001, Student’s t-test).

### Intervention on energy metabolism by metformin prevents cardiac dysfunction in ACHD

Given the newly identified role for *Nkx2-5* regulating metabolism in ACHD hearts, we tested if early pharmacological intervention on global metabolism could prevent heart disease progression in ACHD mice. Young mice under HF diet were treated with metformin, a widely used FDA-approved drug used to treat type 2 diabetes (**Fig 5A**). Metformin acts by improving metabolism, increasing glucose sensitivity, FAO utilization and mitochondrial function (Rena et al, 2017), all processes found dysregulated in ACHD hearts. Both control and ACHD mice treated with metformin showed significant improvement in glucose handling (**Fig 5B**). An early effect in fat mass gain (not shown) was seen in control mice only (15 weeks of age; 9 weeks of treatment), while both control and ACHD mice showed reduced body weight with longer treatment (30 weeks of age; 22 weeks of treatment) (**Fig 5C**). Early differences in weight loss between control and ACHD animals might reflect differential glucose metabolism (**Fig 2B**). As detected earlier, ACHD mice at 15 weeks had a small decrease in EF when compared to control. Short metformin treatment led to increased diastolic and systolic volumes that were independent of genotype (**Fig 5D**, upper panel). At 30 weeks of age (22 weeks of treatment), metformin treatment reverted heart dysfunction on ACHD mice, normalizing EF to control levels by changes in both diastolic and systolic functions when compared to no-metformin ACHD mice (**Fig 5D**, lower panel). These data demonstrate that modulation of energy metabolism by metformin can be potentially used for treatment of cardiac dysfunction mediated by obesity in ACHD.

**Figure 5.**
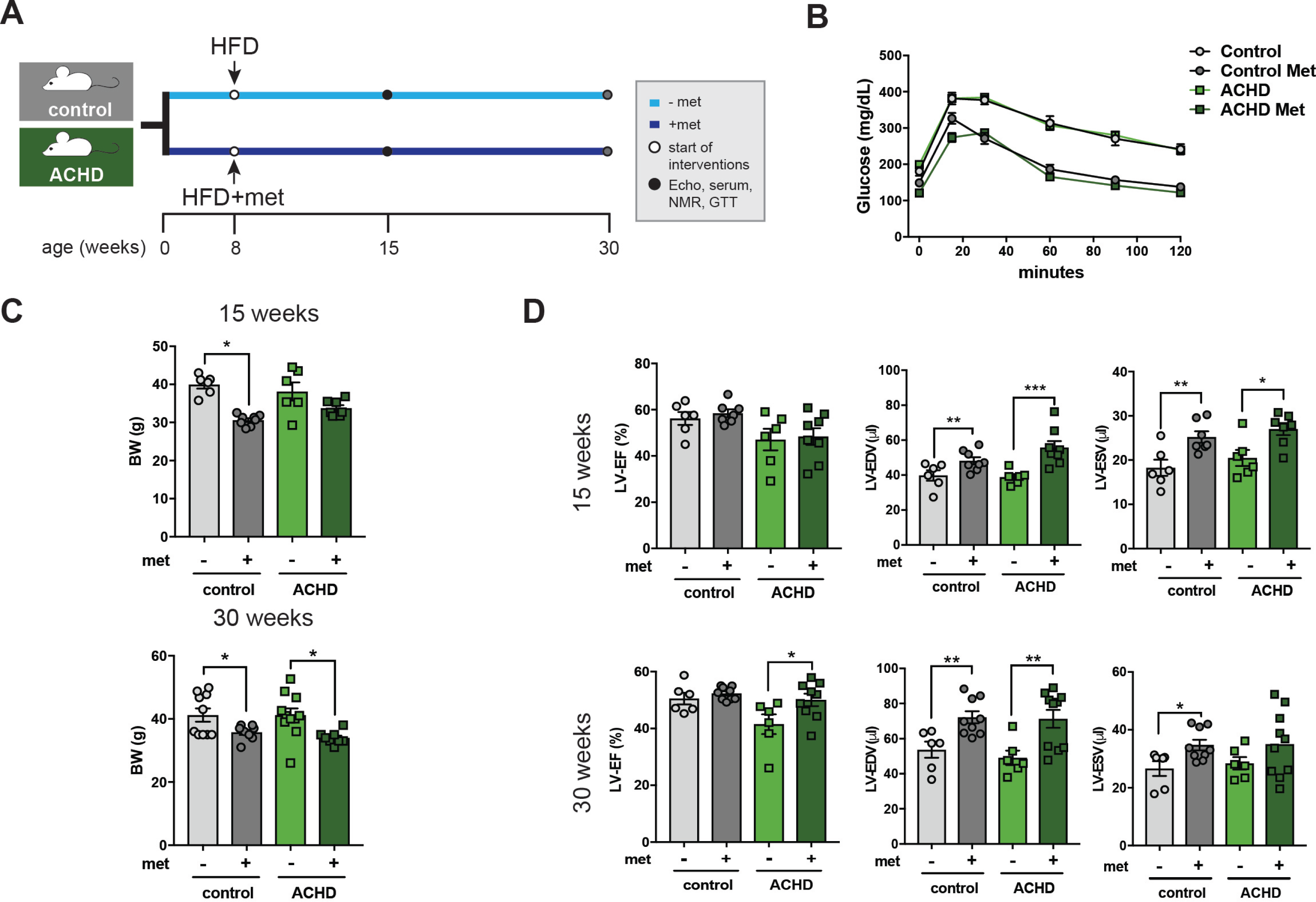
Early metformin treatment prevents cardiac dysfunction in ACHD. (A) Experimental groups and design. (B) Metformin leads to improved glucose handling when compared to controls at 30 weeks of age on both control and ACHD mice. (C) Body weight measurements show that metformin leads to a significant decrease in weight on control mice at 15 and 30 weeks of age while ACHD mice only display decreased body weight at 30 weeks of age. (D) Cardiac functional in control and ACHD mice treated with metformin shows prevention of cardiac dysfunction in ACHD at 30 weeks under HF diet as seen by the significant increase in left ventricle ejection fraction (LV-EF). No significant changes were seen at 15 weeks of age. (N=6; Mean ± SEM, *p<0.05, **p<0.01; ***p<0.001, Student’s t-test). Met: metformin; LV-EF: left ventricular ejection fraction; LV-EDV: left ventricular end diastolic volume; LV-ESV: left ventricular end systolic volume.

## Discussion

Using murine modelling, we demonstrate that changes in energy flexibility is a hallmark of ACHD hearts at early adulthood, and that metabolic imbalance is strongly associated with the progression to heart dysfunction in obesity. Our data show a series of alterations in metabolic substrates that precede cardiac dysfunction in ACHD mice and that pharmacological intervention with metformin can be used to prevent pathological metabolic changes and consequently retain normal heart function. By demonstrating that *Nkx2-5* mutations, known to cause congenital cardiac malformations, are responsible for primary defects in the metabolic capacity of the heart, this study highlights how CHD genes can directly impact cardiac energy utilization. The heart is very sensitive to metabolic imbalance: given the low oxygen and high glucose availability provided by the mother, embryonic and early neonatal hearts display a high dependence on glycolysis. After birth, cardiac metabolism is shifted to fatty acid, which is a more efficient source of energy to sustain the increased demand of the adult heart (Wende et al, 2017). Although >95% of the ATP produced in the adult heart derives from oxidative phosphorylation, energetic flexibility is essential for adaptation of heart function to physiological changes. Decreased flexibility is also associated with pathological states (Ritterhoff & Tian, 2017; Wende et al, 2017), as energy products such as ATP, acetyl-CoA and diacylglycerides directly impact on heart contractility and act as second messengers controlling heart function (Ritterhoff & Tian, 2017).

Given the high energetic demand of the heart and the mitochondrial role in controlling energy consumption, generation of reactive oxygen species and apoptosis, mitochondrial defects are strongly associated with cardiac dysfunction (Huss & Kelly, 2005). The decreased mitochondrial density in ACHD hearts suggests a primary defect in mitochondrial biogenesis, in which further investigation is merited. Mitochondrial dysfunction is also supported by the observed decrease in cellular respiration and impaired energetics in ACHD hearts, where early changes in cardiac metabolism lead to increased dependency on glycolysis and decreased metabolic flexibility. Altered glucose metabolism is one of the drivers of the hypertrophic response and an early marker of heart disease progression (Kundu et al, 2015), which could be adaptive under a homeostatic condition, but triggers heart dysfunction once a second metabolic stress (such as obesity) is imposed.

Despite the extensive energetic abnormalities in ACHD hearts, no evidence of lipotoxicity was found in histological analyses of heart sections (not shown), although important changes in lipid distribution were detected in the metabolomic analyses. Given the decreased energetic flexibility and fatty acid oxidation coupled with increased fatty acid availability in ACHD/obesity, it is possible that lipotoxicity may play a role in the later stages of disease progression.

Metformin has cardioprotective effects and limits cardiovascular events in patients (Eppinga et al, 2017; Lexis et al, 2014), regulating global and tissue metabolism via complex mechanisms that are not completely understood. Metformin acts by reducing hepatic glucose production by the gut and by increasing glucose organ sensitivity (Pernicova & Korbonits, 2014). At the cellular level, metformin likely acts via AMPK-dependent and –independent mechanisms to modulate mitochondrial and lysosome function (Rena et al, 2017). Interestingly, despite the early detection of high levels of ATP in ACHD hearts at 8 weeks, no changes in AMPK were detected, suggesting a possible impairment of the AMPK signalling pathway, or an AMPK-independent mechanism of action (**Fig 4G, H**). Recent studies have shown that chronic exposure to metformin increases lifespan in mice (Martin-Montalvo et al, 2013; Valencia et al, 2017), has beneficial effects in cancer treatments (Heckman-Stoddard et al, 2017) and cardiovascular dysfunction (Eppinga et al, 2017; Griffin et al, 2017; Kobashigawa et al, 2014; Sun & Yang, 2017; Tzanavari et al, 2016), albeit some divergence between preclinical and clinical trials for metformin in humans have been described (Lexis et al, 2014). as seen for non-diabetic STEMI patients. Cellular uptake of metformin is controlled by organic cation transporters (OCT) belonging to the *Slc22a* gene family. Expression of OCT1 and OCT3 have been described in the heart (Lozano et al, 2018). Future studies are necessary to establish the tissue-specific action of global metformin administration to mice and cardiac-intrinsic mechanisms determining beneficial effects of metformin treatment in ACHD.

## Conclusions

The increased prevalence of ACHD in the population (Alshawabkeh & Opotowsky, 2016; Gilboa et al, 2016; Marelli et al, 2014) and its associated with metabolic syndrome (Deen et al, 2016) calls for mechanistic studies to understand how cardiac energy flexibility correlates with heart function in homeostasis and disease (Goodpaster & Sparks, 2017; Neglia et al, 2007; Ritterhoff & Tian, 2017). Surveillance of cardiac performance combined with modulation of energy utilization could lead to improvement of heart function and prevent heart failure in ACHD, as demonstrated in this study using murine modeling. More effective treatments for heart failure should move focus from solely examining neurohormonal/unloading modulation to including corrections for metabolically affected hearts (Brown et al, 2017). The findings presented here argue for manipulation of energy metabolism and mitochondrial function as potential intervention targets to revert progression to heart failure in ACHD.

## Acknowledgments

We thank Pete A. Williams for providing mitochondrial antibodies and advice, and the *In Vivo* Physiology Core of the Jackson Laboratory for the generation of all metabolic and physiological data.

## Author Contributions

Conceptualization, M.W.C and M.B.F.; Methodology, J.C.W., N.A.R., M.B.F. and M.W.C.; Formal Analysis, G.C., P.B., V.P., S.A. and M.R.; Investigation, J.C.W., R.P., A.C., O.H., J.S., Q.W., J.T.P, M.B.F. and M.W.C.; Writing – original draft, J.C.W., M.B.F. and M.W.C.; Writing – Review and Editing, J.H., H.P. and N.A.R.; Funding Acquisition, N.A.R. and M.W.C.; Supervision, M.B.F., J.H., N.A.R., M.W.C.

---

**Supplementary Figure 1.**
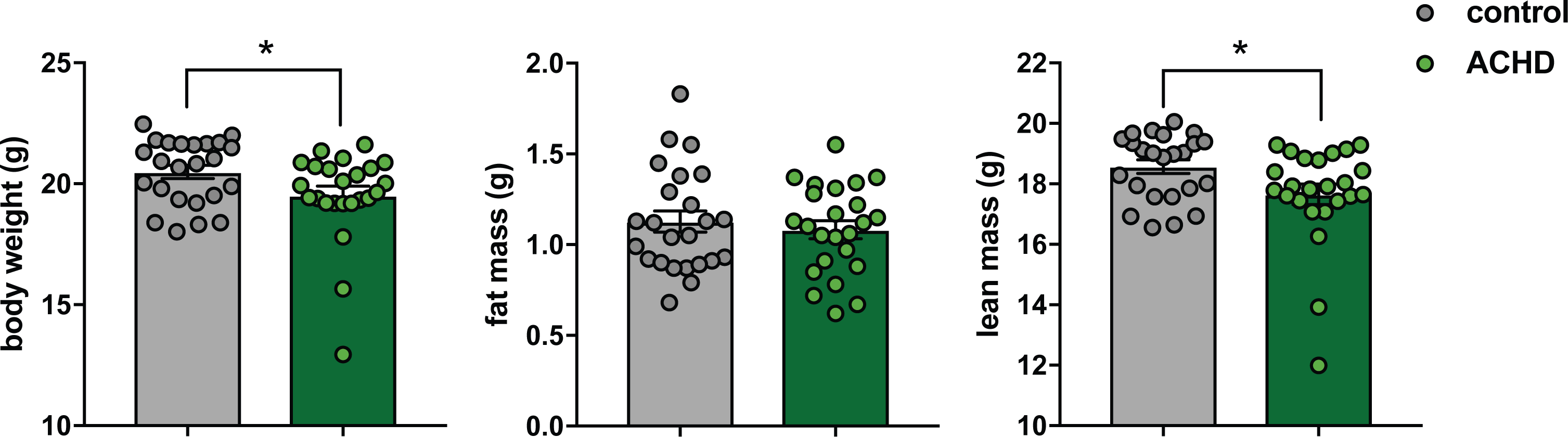
Decreased body weight and lean mass in ACHD mice.

**Supplementary Figure 2.**
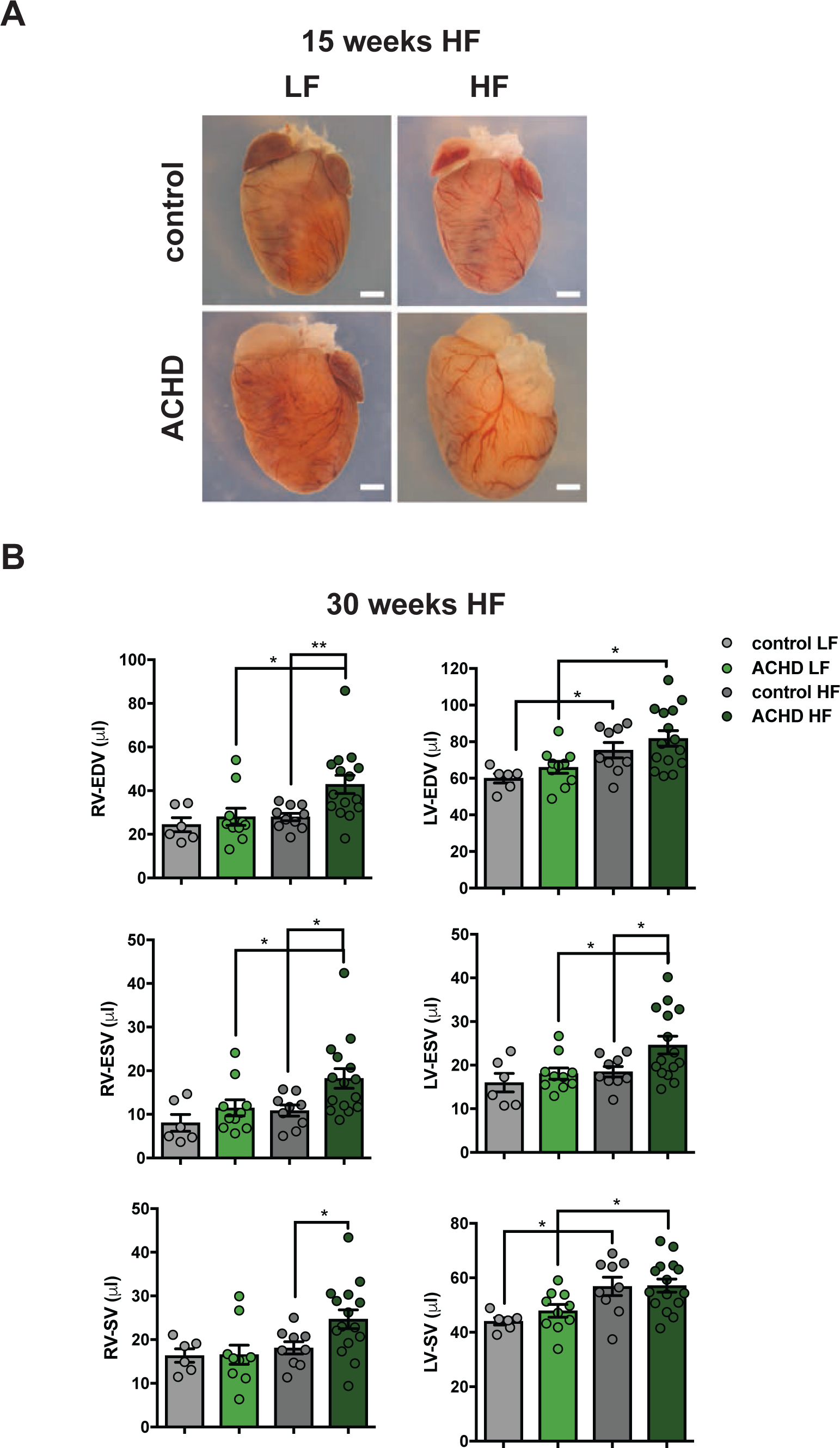
Analysis of ACHD aged mice. (A) Whole mount hearts at 15 weeks of age where no significant change in cardiac function is detected. (B) Analysis of heart function by MRI on 30-week old mice show significant right ventricle (RV) and left ventricle (LV) dysfunction that is associated with dilation (increased end-diastolic and systolic volumes in ACHD HF mice. EDV-end diastolic volume; ESV-end diastolic volume; SV-stroke volume. (Mean ± SEM, *p<0.05, **p<0.01, Student’s t-test).

**Supplementary Figure 3.**
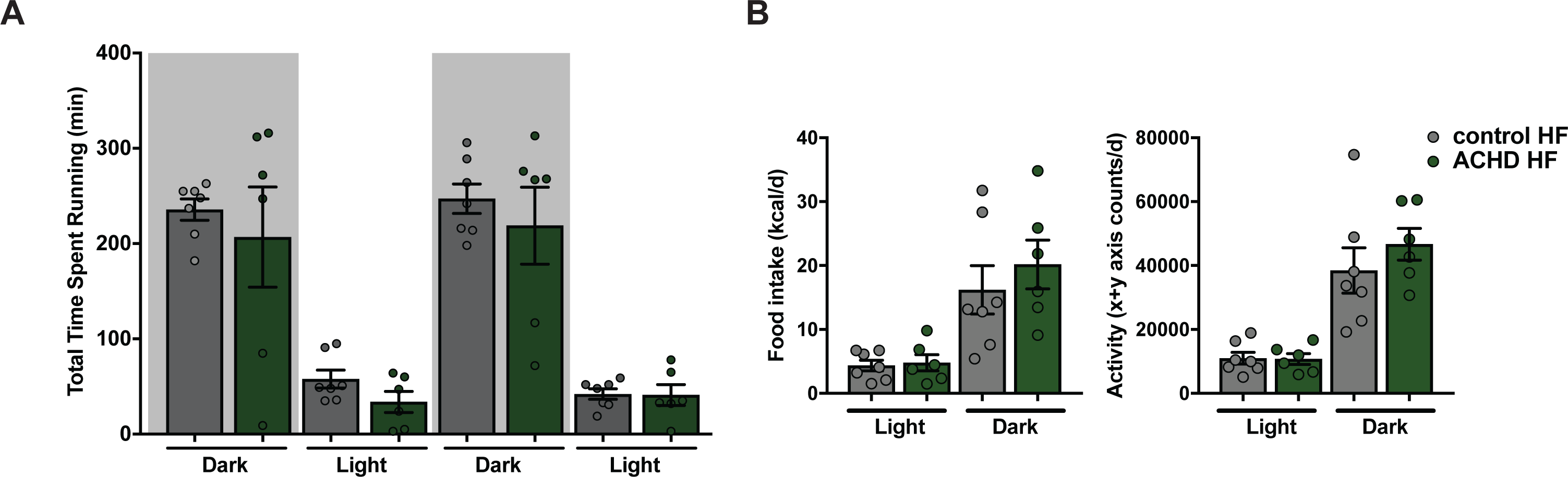
Additional physiological parameters in ACHD mice at 30 weeks. Mice on HF diet show (A) no changes in time spent at running wheel under voluntary exercise between control and ACHD mice and (B) no significant difference in food intake or activity.

**Supplementary Figure 4.**
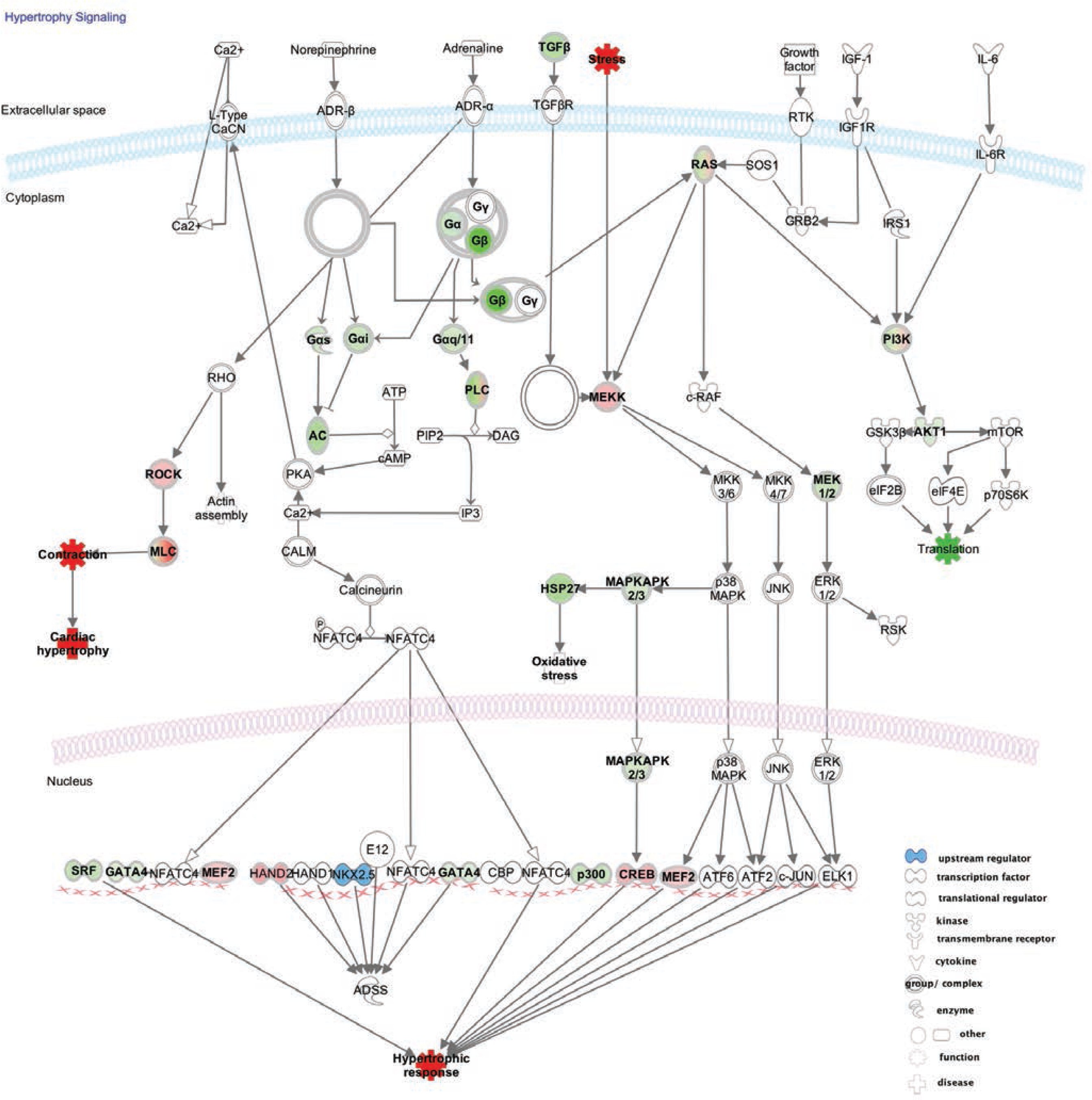
Predicted changes in hypertrophic signalling seen in adult ACHD hearts using Ingenuity Pathway Analysis. Green: down-regulated genes; red: up-regulated genes. Nkx2-5 is shown in blue.

**Supplementary Figure 5.**
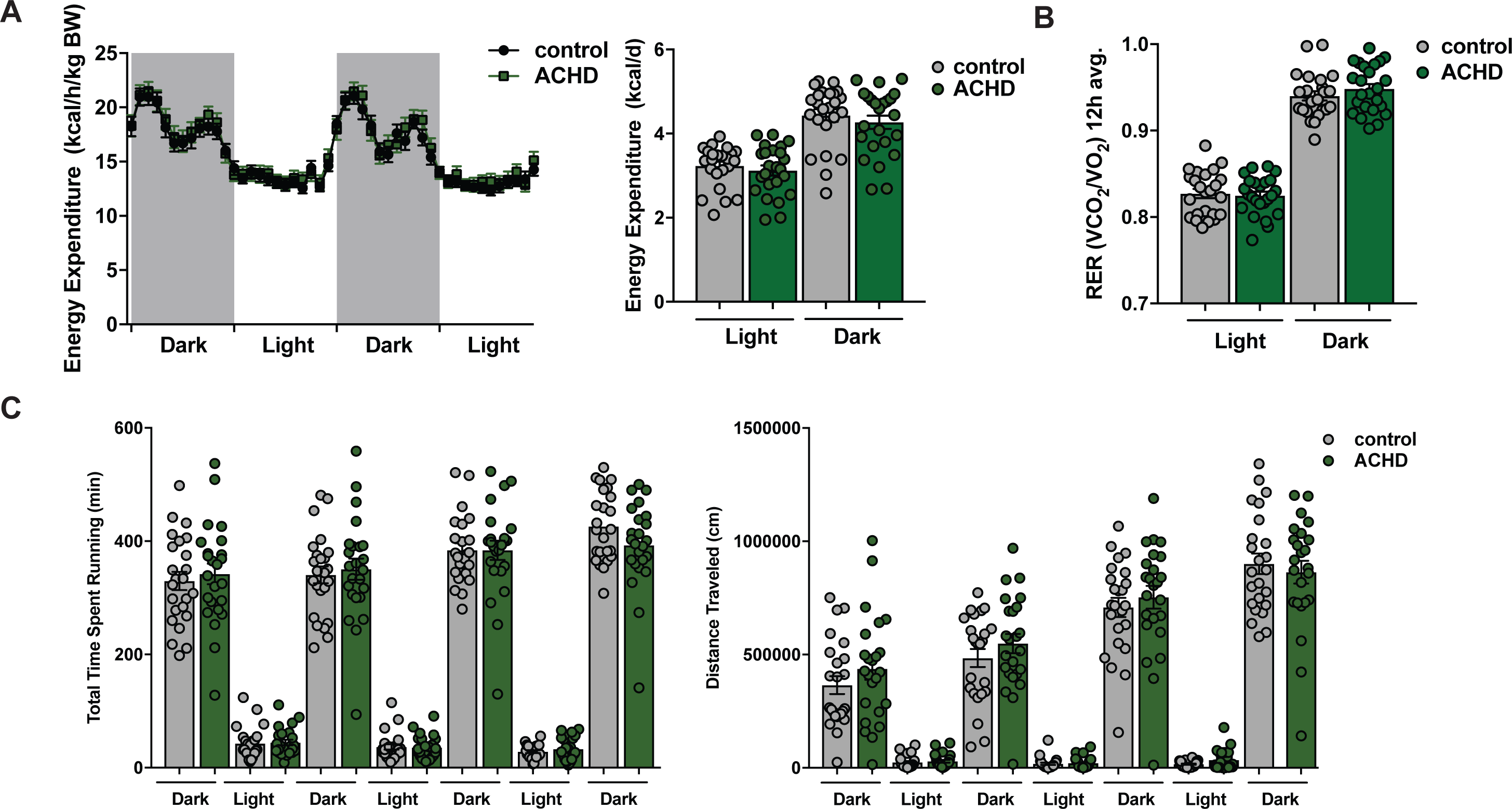
Metabolic changes in ACHD mice at 8 weeks. (A-C) No significant changes were seen in energy expenditure, RER or voluntary running capacity between control and ACHD mice.

**Supplementary Figure 6.**
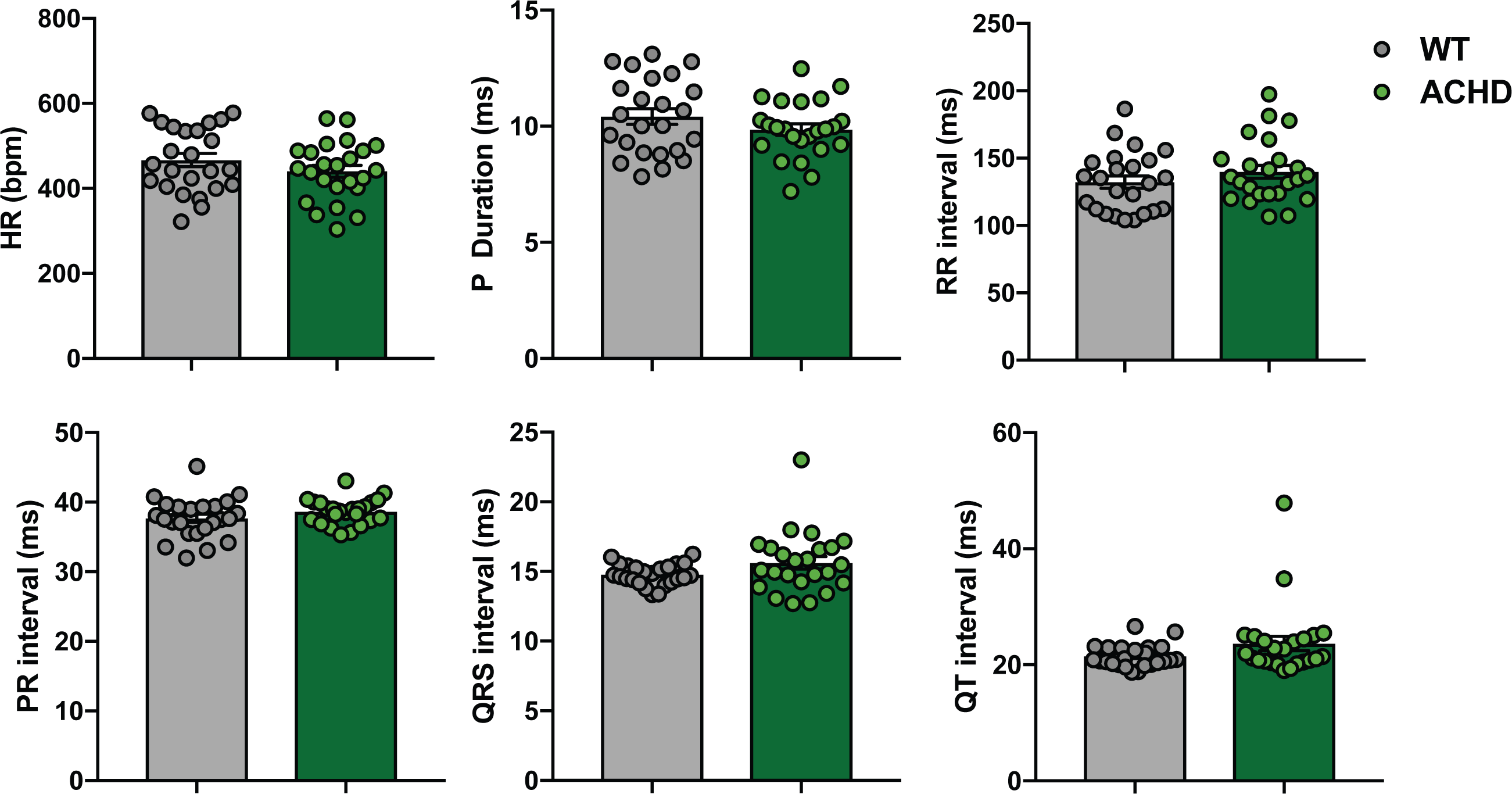
ECG analysis. No changes in electrical profiling were detected between control and ACHD mice at 8 weeks of age.

**Supplementary Table 1.**
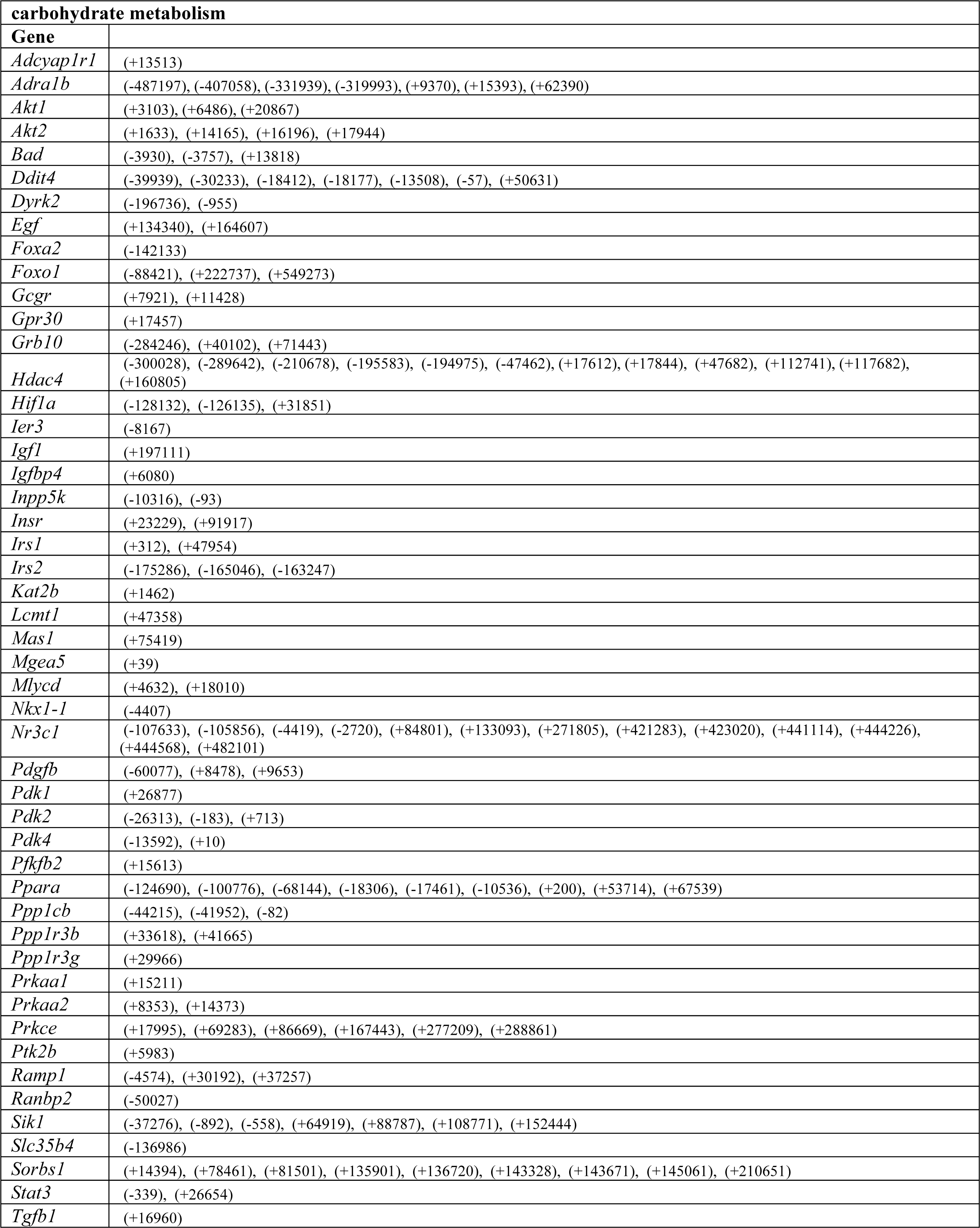
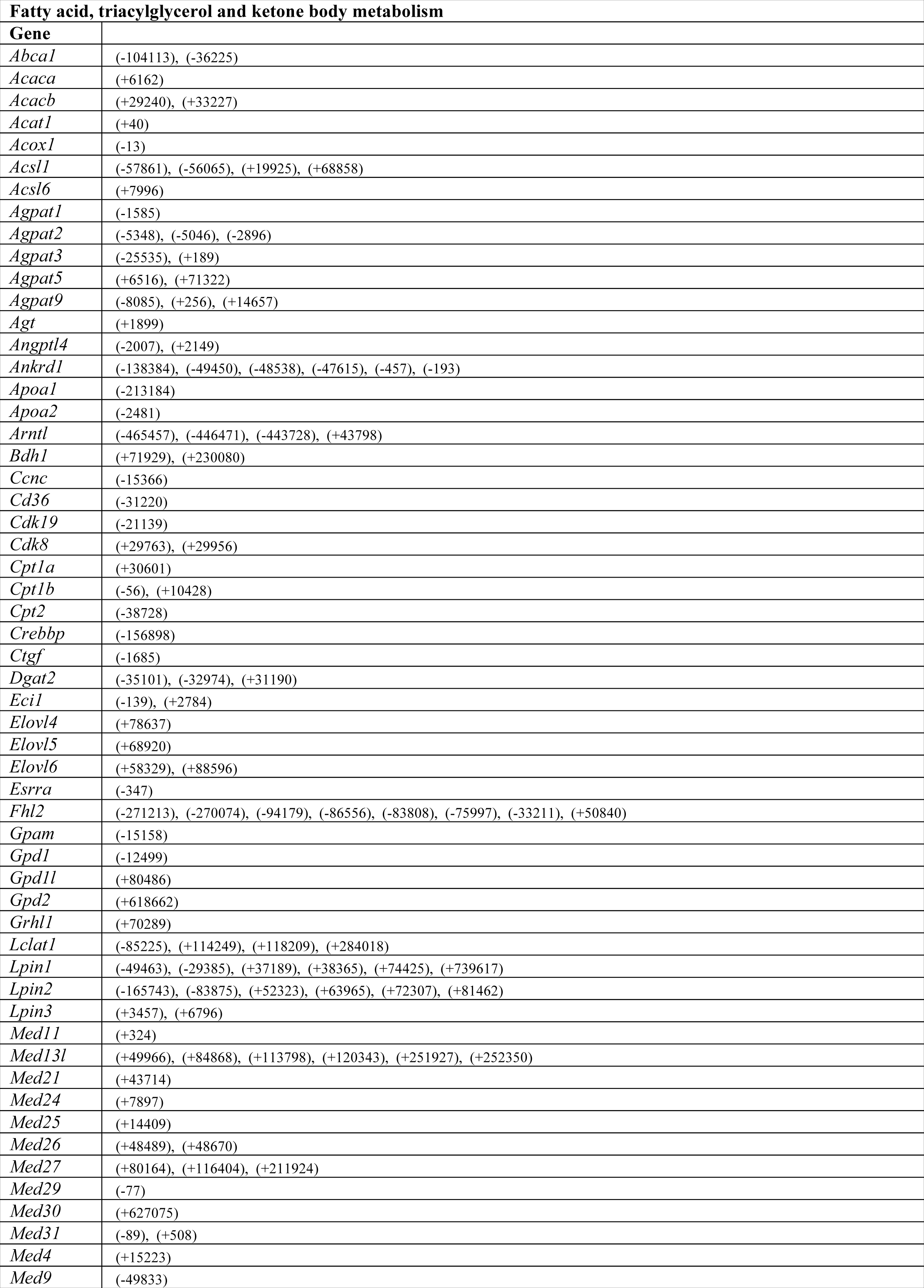
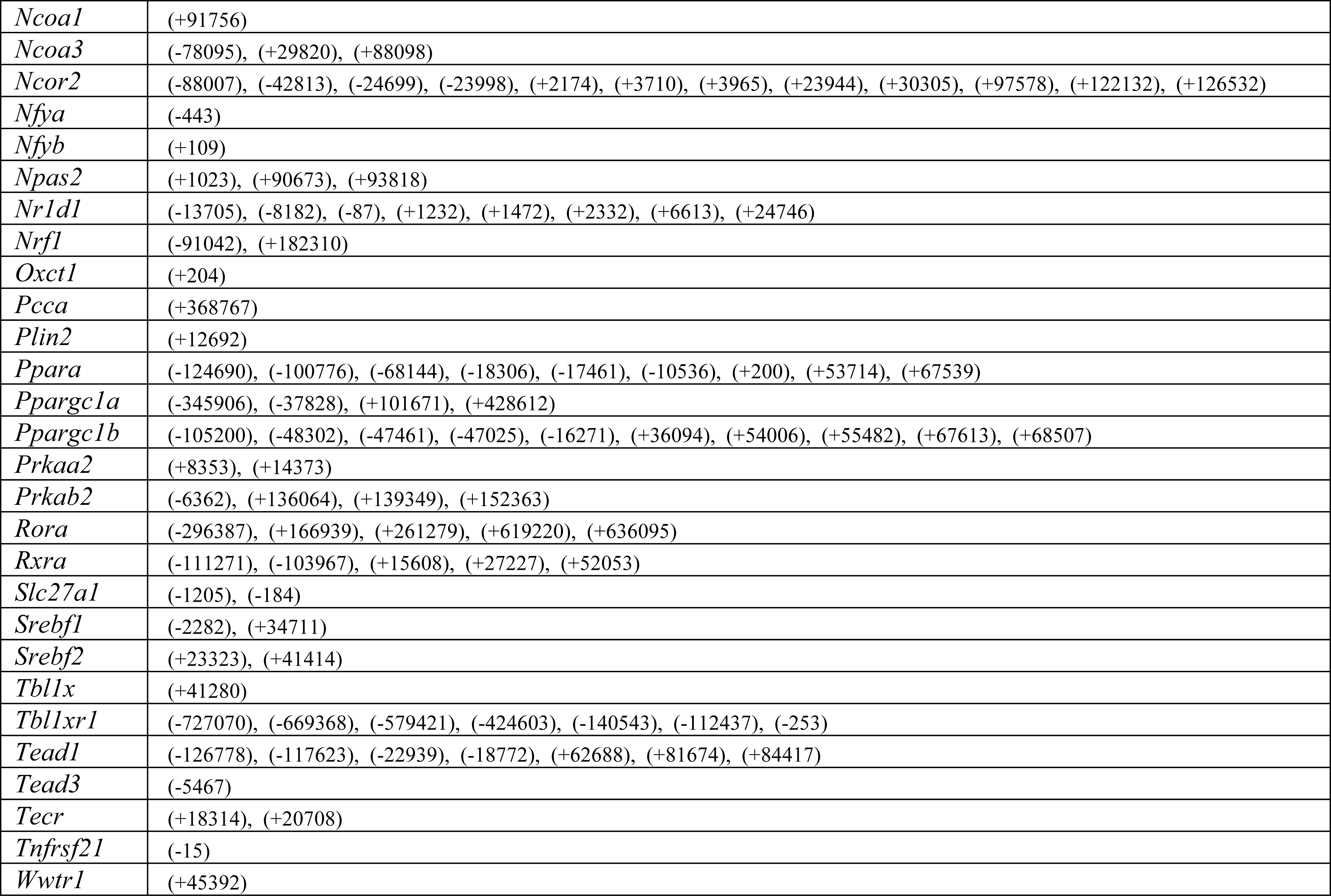
ChIPseq analysis of adult heart show high prevalence of genes associated with metabolic processes.

**Supplementary Table 2.** Echocardiographic analysis of ACHD hearts at 8 weeks.

| | EF (%) | FS (%) | LV Mass mg | LV Vol;d μl | LV Vol;s μl | IVS;d mm | IVS;s mm | LVID;d mm | LVID;s mm | LVPW;d mm | LVPW;s mm | N |
| --- | --- | --- | --- | --- | --- | --- | --- | --- | --- | --- | --- | --- |
| **control** | 48.69±9.78 | 24.40±6.20 | 92.6±20.14 | 61.43±11.82 | 31.73±8.38 | 0.77±0.12 | 0.92±0.09 | 3.76±0.32 | 2.85±0.36 | 0.72±0.09 | 0.90±0.11 | 24 |
| **ACHD** | 46.27±8.67 | 22.95±5.24 | 100.0±22.7 | 68.06±12.16 | 36.91±10.12 | 0.71±0.09* | 0.88±0.14 | 3.96±0.28* | 3.07±0.31* | 0.73±0.16 | 0.90±0.19 | 24 |

**Supplementary Table 3.**
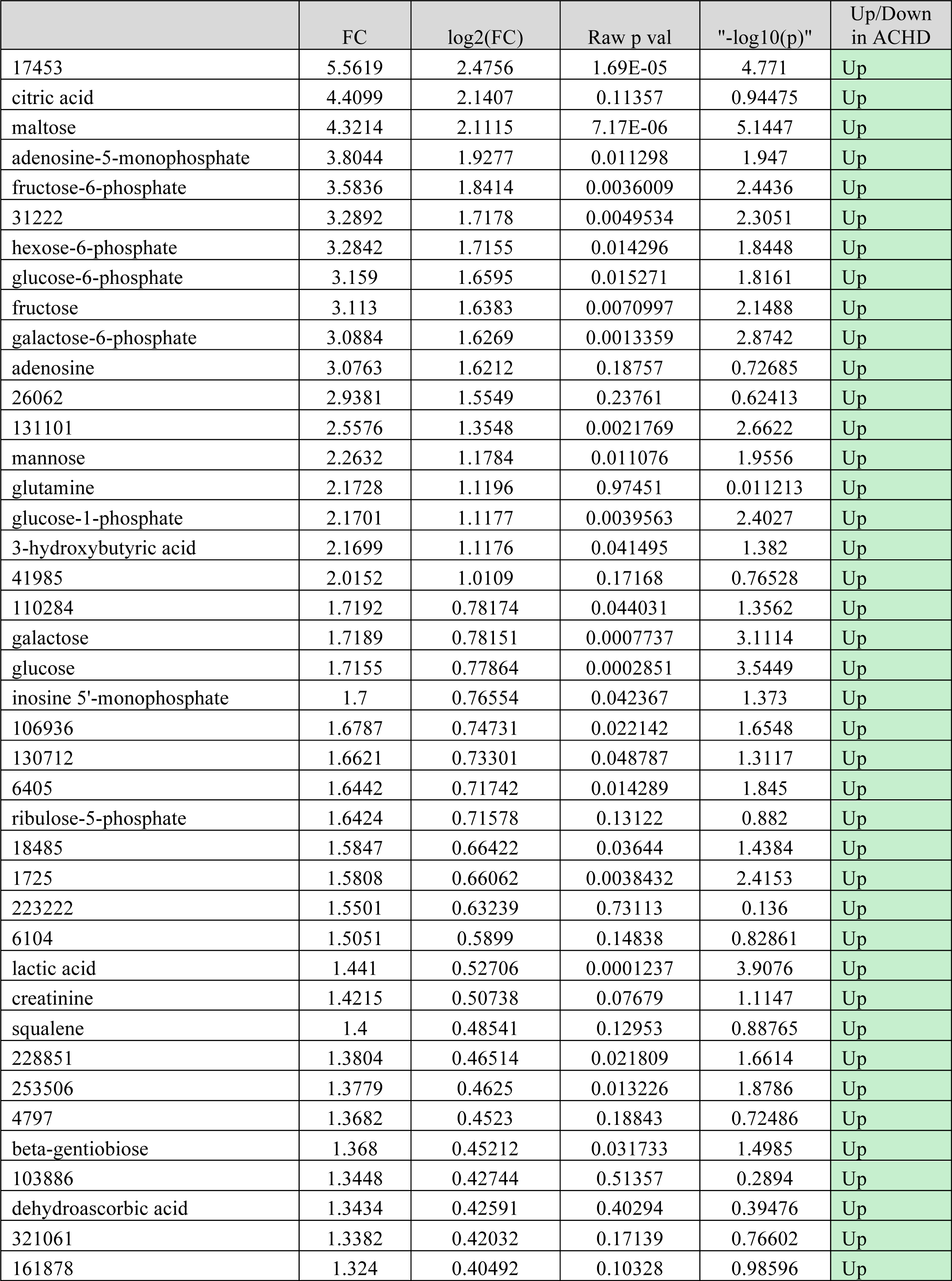
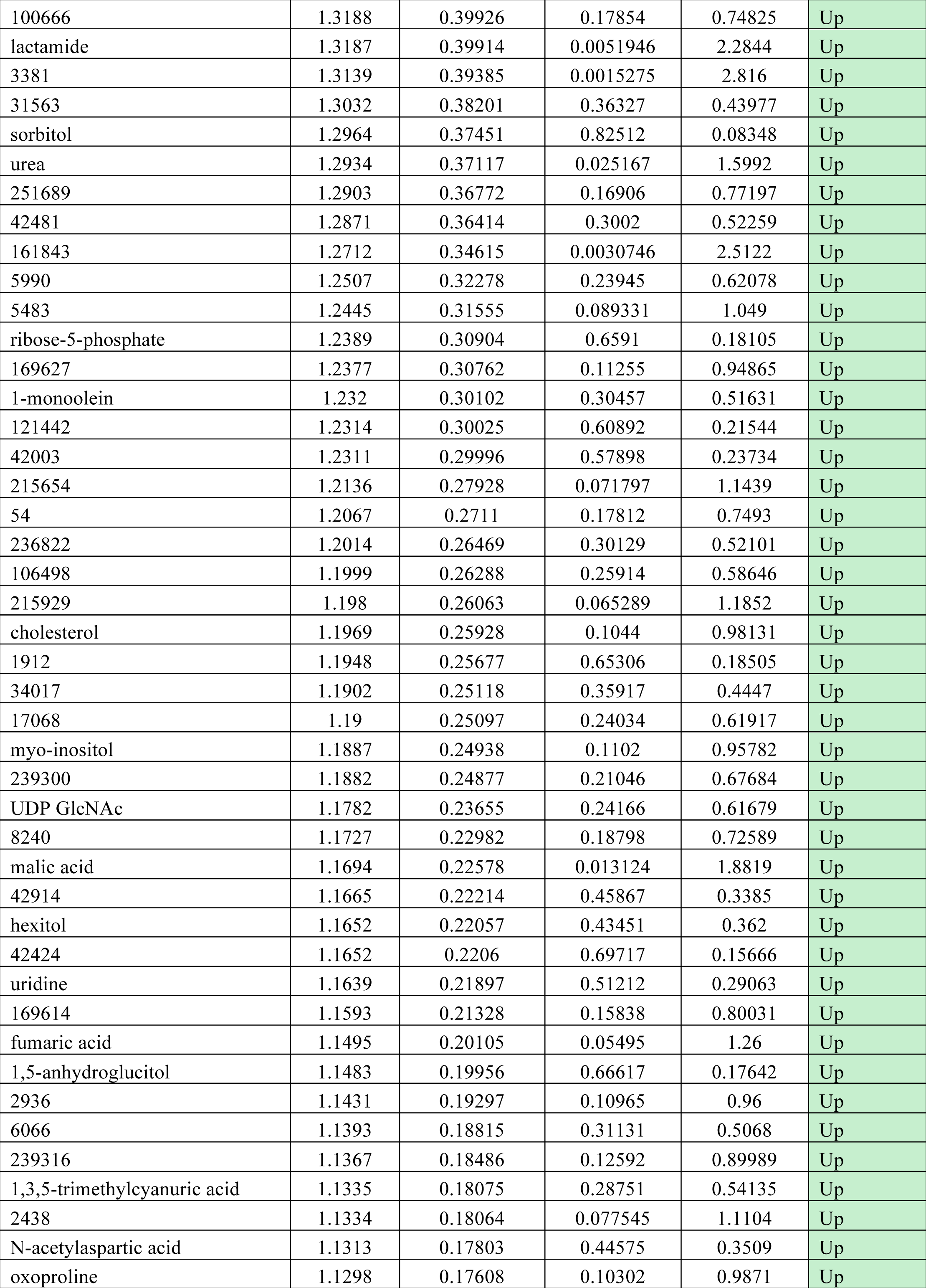
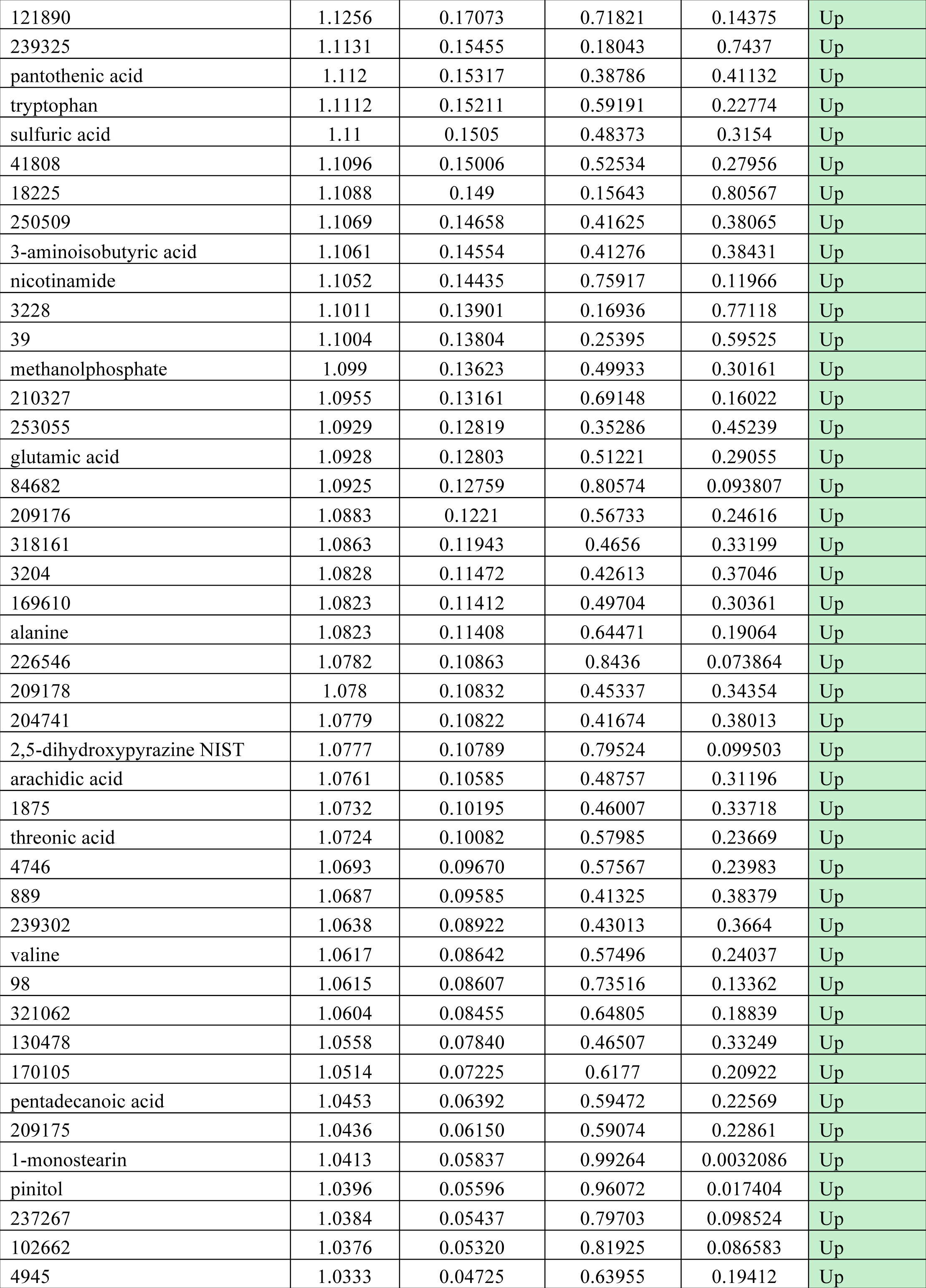
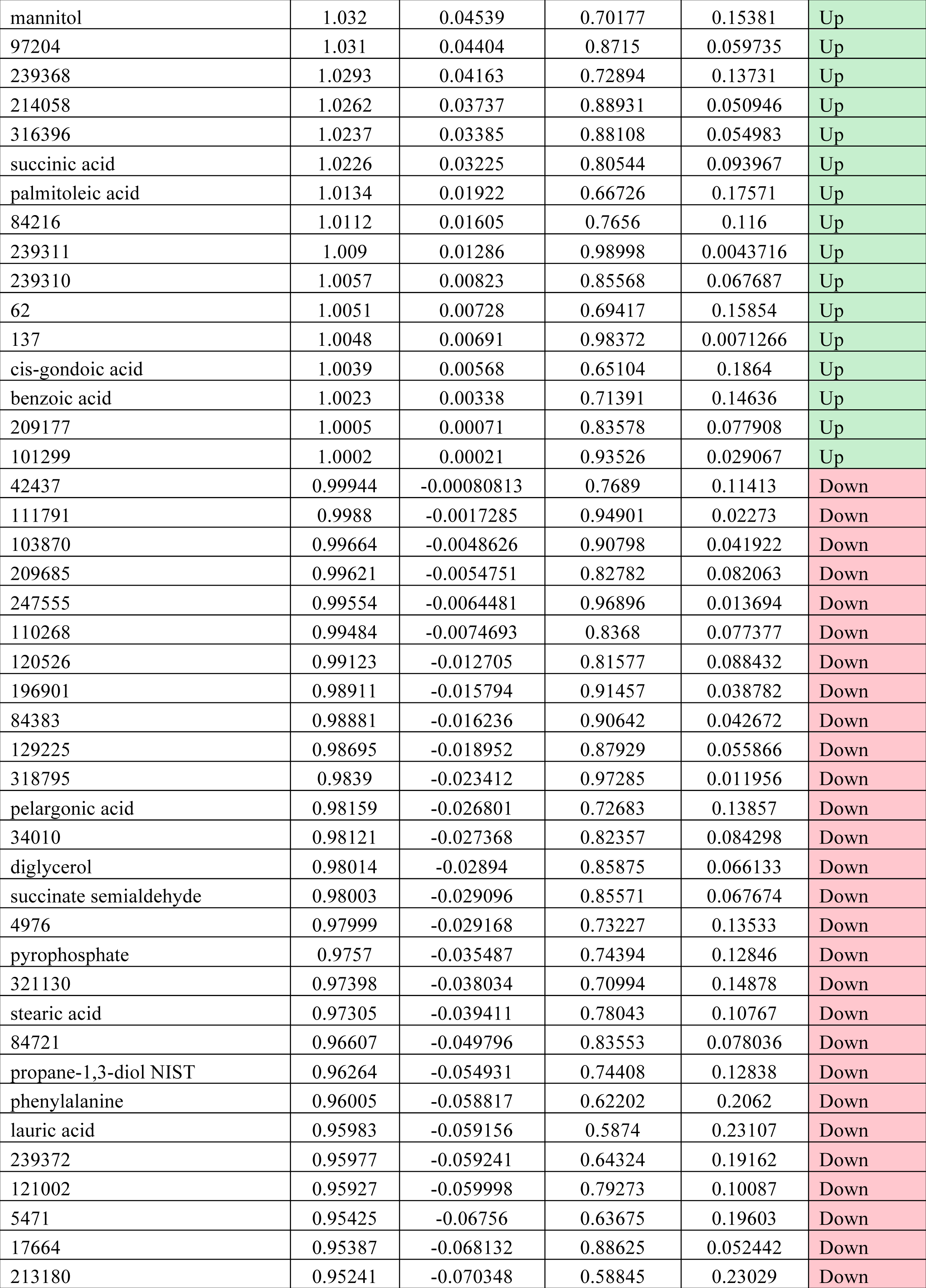
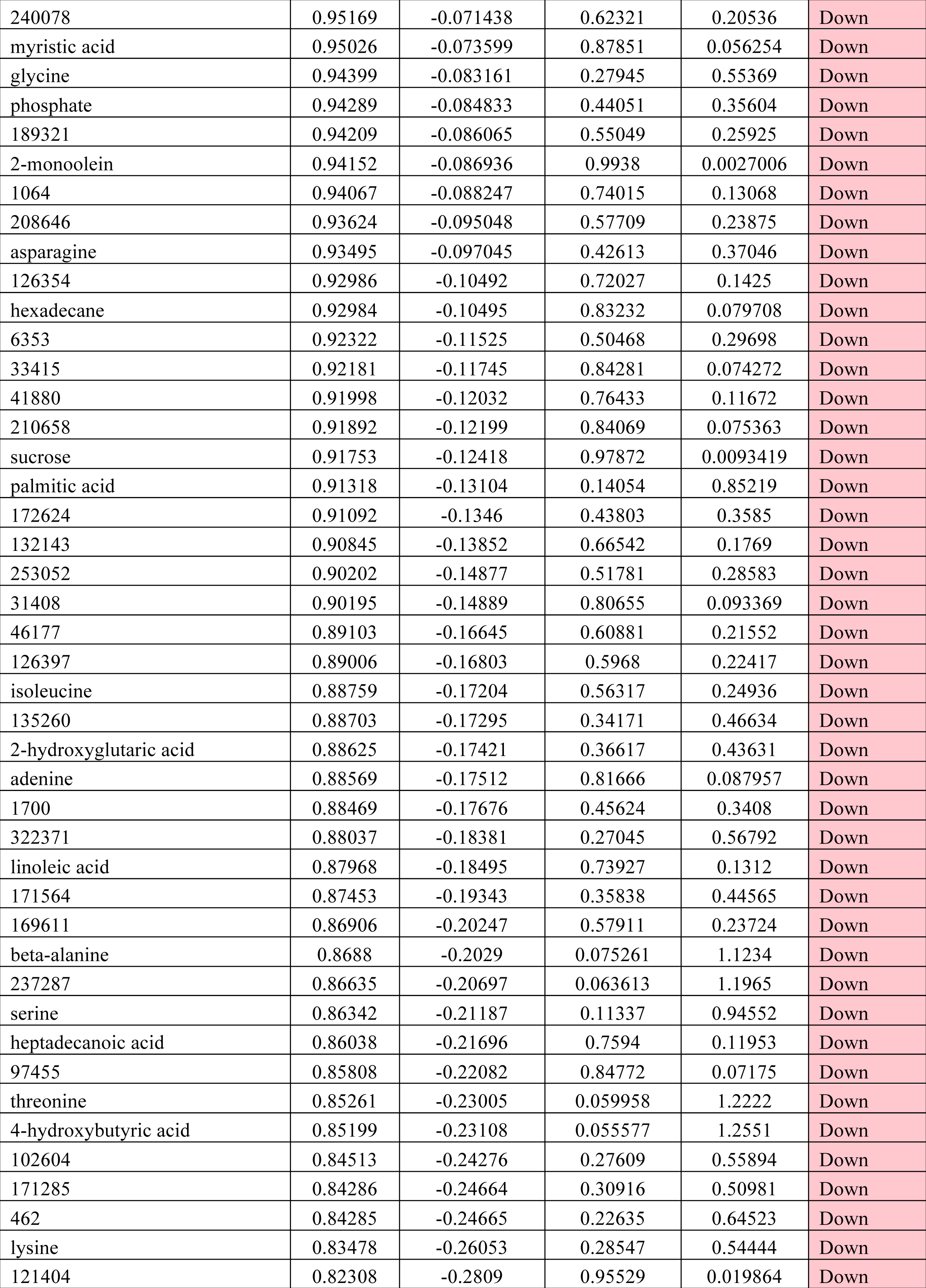
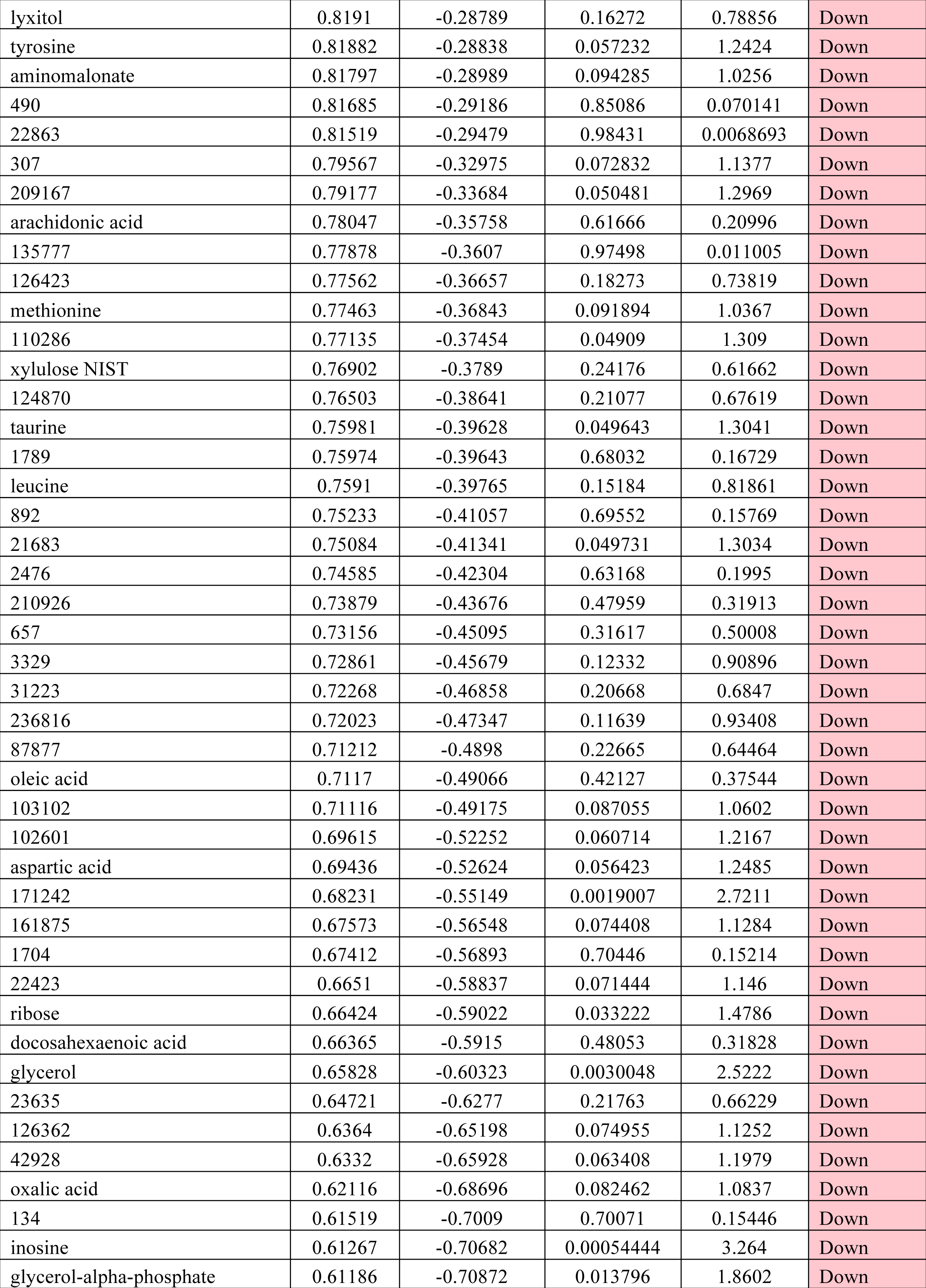
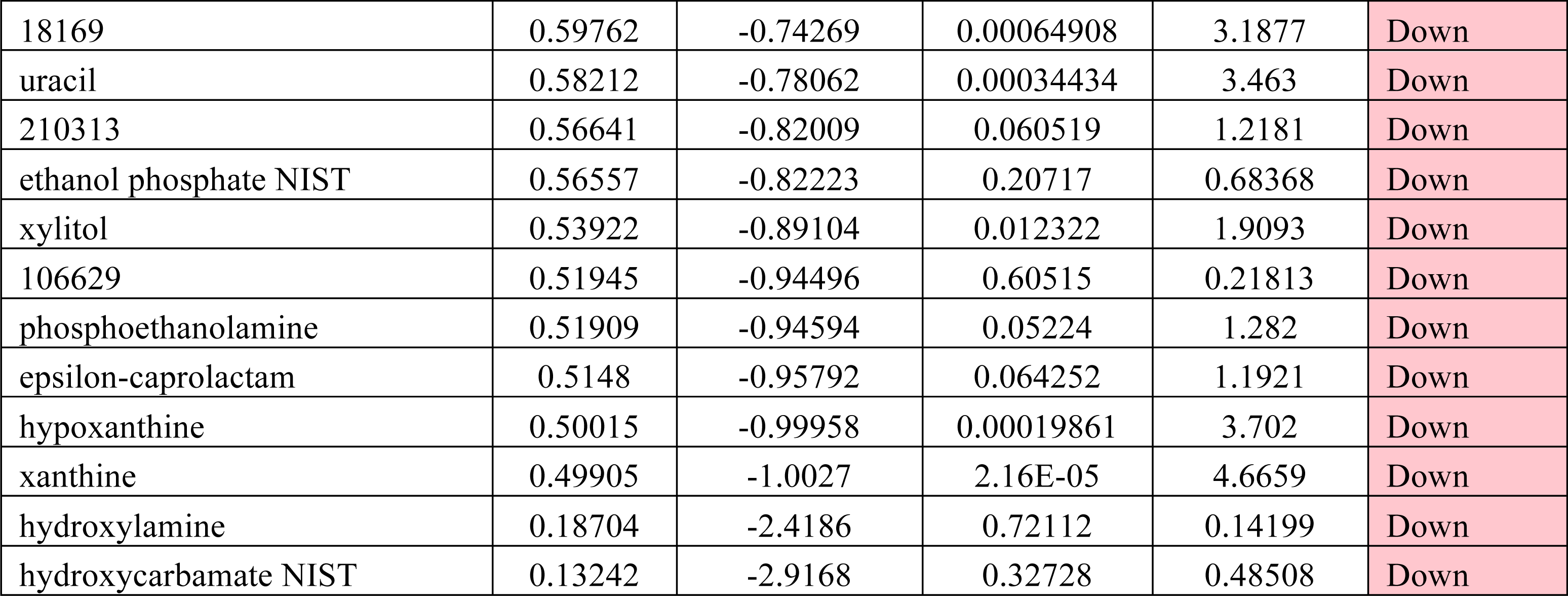
Changes in metabolites seen in ACHD hearts at 8 weeks.

**Supplementary Table 4.**
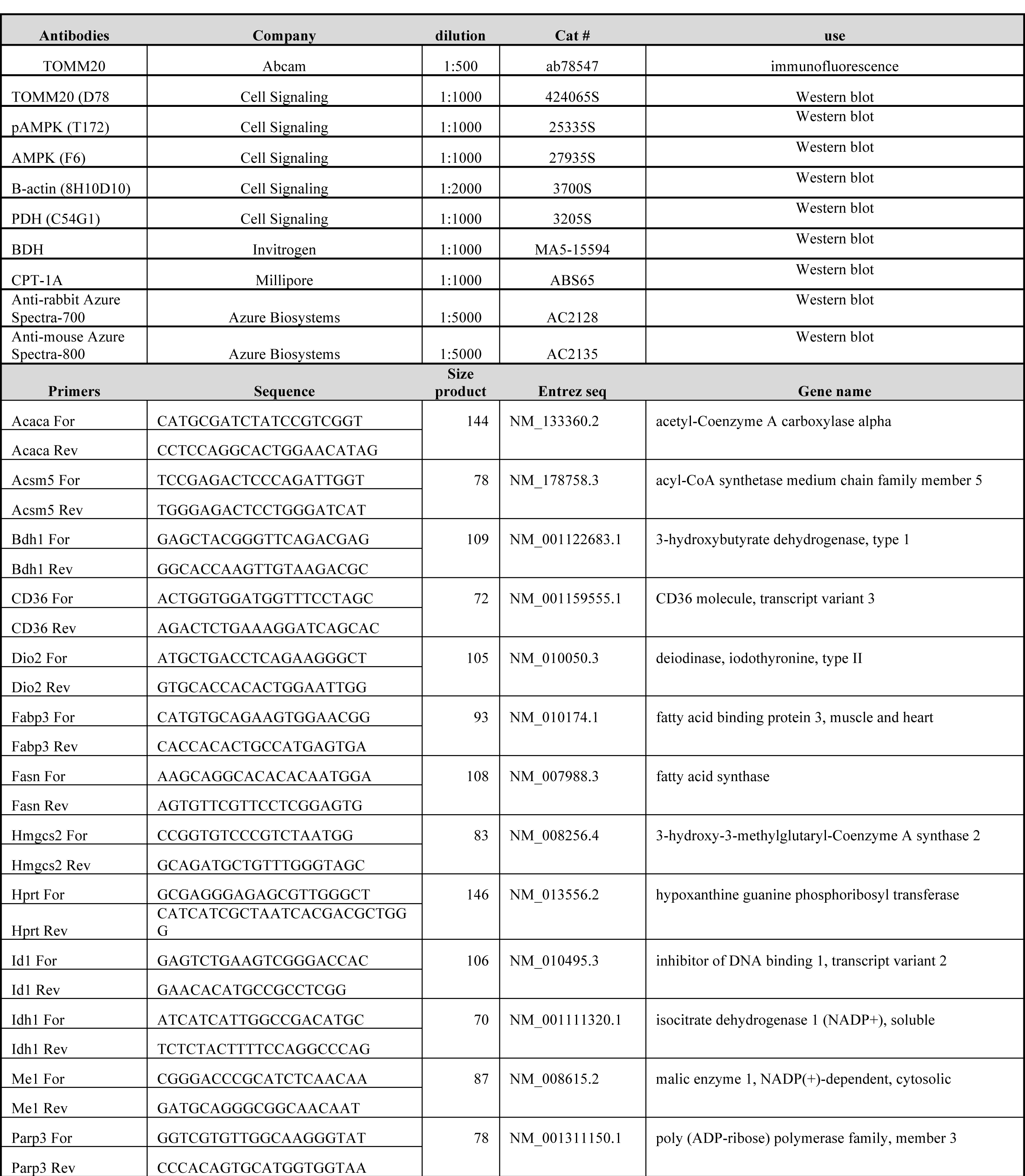
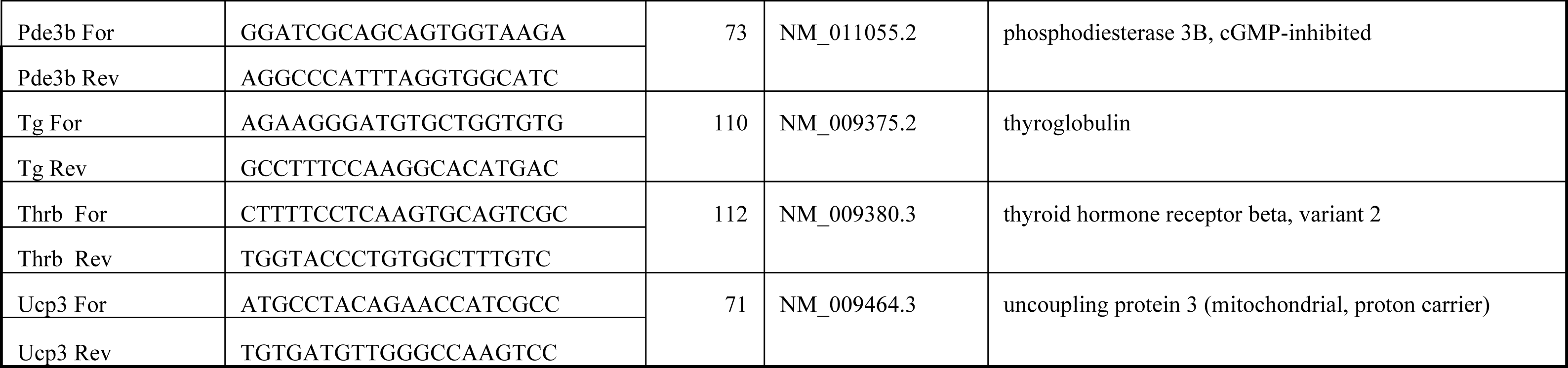
Antibodies and Primers used

